# A non-canonical role for the ATAD2 ortholog BDF7 in nucleolar ribosome maturation and amastigote survival in *Leishmania mexicana*

**DOI:** 10.64898/2025.12.23.692120

**Authors:** Nathaniel G. Jones, Juliana Brambilla Carnielli Trindade, Alexander Johnson, Adam Dowle, Raquel Gabarró, Félix Calderón, Anthony J. Wilkinson, Jeremy C. Mottram

**Author notes:** ViiV Healthcare, 36 E Industrial Rd, Branford, CT 06405, United States.

## Abstract

ATAD2 is a widely conserved, homohexameric ATPase that, in humans, features a bromodomain tier. It plays a role in chromatin remodelling in embryonic stem and germ cells but is frequently upregulated in many cancers, making its bromodomain an attractive drug target. While ATAD2-like proteins generally modulate nucleosome density to control genome compartmentalization and gene expression across eukaryotes, their precise molecular functions vary. We investigated LmxBDF7, the ATAD2 ortholog in the important human pathogen *Leishmania mexicana*, the causative agent of cutaneous leishmaniasis. Unlike its human counterpart, the LmxBDF7 bromodomain is predicted to be occluded and non-canonical. BDF7 null mutants were unable to develop into infectious amastigote forms and could not infect macrophages, demonstrating it is essential for lifecycle progression. Chromatin Immunoprecipitation sequencing (ChIP-seq) suggested low affinity for chromatin, aligning with its atypical bromodomain. Instead, Proximity-Biotinylation (XL-BioID) suggested a role in ribosome maturation within the nucleolus. RNA-sequencing (RNA-seq) of the Δ*bdf7* mutant revealed widespread disruption of gene expression during growth and differentiation. Crucially, key genes required for amastigote survival, such as ribosomal protein genes and glutamine synthetase, were downregulated. This downregulation was spatially biased, preferentially affecting genes on Chromosome 23. Our combined data suggest that while BDF7 is essential, its functions have diverged from other ATAD2-like factors found in opisthokonts.

**Author Summary:** ATAD2 is a protein that helps cells balance the number of nucleosomes bound to DNA in the nucleus of a cell. Occurring at important sites or times this provides a helper function that ensures other protein complexes can operate on chromatin effectively. In some cancers ATAD2 is disrupted and therefore is being explored as target for new medicines. Orthologues of ATAD2 have been characterised in mammals and several species of yeast. We have sought to identify if an ATAD2-like protein can be found in the important human pathogen, *Leishmania mexicana –* which is evolutionary distant from humans and yeast. Indeed, we were able to find an ATAD2-like protein called BDF7. Interestingly, BDF7 has a bromodomain which is predicted to be non-functional in terms of being able to bind histones. We were able to make strains of *Leishmania mexicana* that lacked BDF7 which were viable, but unable to complete the differentiation step required for infecting macrophages. Intriguingly we found BDF7 had a poor association with chromatin and inhabited a protein neighbourhood defined by factors which facilitate ribosome biogenesis. Lastly, RNA-seq analysis of the cells revealed that those lacking BDF7 were potentially depleted for glutamine synthetase which would prevent them developing into fully functional amastigotes.

## Introduction

Parasites in the genus *Leishmania* are important human and animal pathogens, causing a variety of diseases collectively termed leishmaniases ^1^. Ranging from cosmetically damaging ulcers to life-threatening infections of the visceral organs, an estimated 1 million people per year are affected. While several drugs are available to treat leishmaniasis all have negative side effects and an increasing risk of low efficacy^2^ . A pipeline of new chemical entities is undergoing preclinical and clinical development but several of these have been paused due to detection of potential toxicity issues (https://dndi.org/research-development/portfolio/), underscoring the need to maintain active prioritisation of other potential targets. Currently, no vaccine is licensed for use in humans.

*Leishmania* are transmitted to humans by the bite of an infected sand fly and upon which the forms (mainly metacyclic promastigotes, although haptomonad stages may form an important component^3^ ) introduced to the bite-site must be ingested by phagocytic cells for the infection to progress. In these cells the metacyclic form must differentiate into the amastigote form - a process associated with dramatic morphological and metabolic change. *Leishmania* are in the class of kinetoplastids, and clade of discoba, as such they evolved from a deeply rooted branch of eukaryotes. Their study can provide information on the evolution of eukaryotes, as recent analyses suggest the last eukaryotic common ancestor was a cell with excavate features ^4,5^.

We have recently focused on understanding how *Leishmania* parasites use a class of proteins called bromodomains factors (BDFs). These are proteins that use a simple interaction module, the bromodomain, to bind acetylated lysine residues on histones in nucleosomes^6^; the protein hubs that package DNA and act as a platform for regulatory post-translational modifications^7^. Dysregulation of bromodomain function has been associated with disease in particular various cancers, and bromodomains have been explored as potential drug targets, with particular focus on the BET (bromodomain extra terminal) family of bromodomains^8^. Due to toxicity problems with BET inhibitors, researchers are now exploring other classes of bromodomains. One such BDF, ATAD2 (ATPase family AAA domain-containing 2), has been identified as an upregulated factor in a variety of human cancers, and these elevated expression levels correlate negatively with disease prognosis in breast and lung cancers ^9,10^ . As the bromodomain is a small interaction module, BDFs must exert their action on chromatin using accessory domains or protein partners. ATAD2 has a tandem arrangement of AAA+ ATPase domains (‘ATPases associated with diverse cellular activities’), one active and one inactive, followed by a bromodomain in the C-terminal region of the protein. ATAD2, like other AAA+ ATPases forms ^11^ a homohexameric complex with a central pore that appears to act as the substrate binding site^12^. The ATPase domains of ATAD2 and related proteins form a 2-tiered ATPase assembly arranged in a shallow spiral with a third tier of bromodomains atop the active ATPase ring. Nucleotide and substrate binding cause rearrangements in the structure of this ring and pull substrates into the pore, with these structural studies focusing on the H3/H4 dimer ^12–14^.

In mammals, ATAD2 has been reported to be almost exclusively expressed in embryonic stem (ES) cells and male germ cells^15^ with its expression in malignant cells and tumours being correlated with pathology ^9^. Under normal physiological conditions in ES cells, ATAD2 is recruited to acetylated histones and ensures the dynamic nature of chromatin by promoting a high rate of histone exchange^15^. This is conducted by ensuring the turnover of the histone chaperones HIRA and FACT complexes^16^. ATAD2 is not essential for survival, as ATAD2 knockout mice can be generated. Using this mouse model, ATAD2 has recently been reported to influence HIRA-H3.3 dependent histone to protamine exchange during spermatogenesis, resulting in poorly compacted genomes in the sperm cells ^17^.

ATAD2-like proteins are readily identifiable in non-human animals and other members of opisthokonta such as yeast, and are suggested to be well conserved in evolution^18,19^ . However, most work on ATAD2 and its homologues has focused on the enzymes from opisthokonts such as human (*Homo sapiens*), mouse (*Mus musculus*) and brewer’s yeast (*Saccharomyces cerevisiae*). The *S. cerevisiae* Yta7 protein for example has been reported to maintain the balance of nucleosomes in chromatin – its disruption results in changes to histone gene transcription and DNA replication ^20–25^. Yta7 has been shown to act as an S-phase cyclin-dependent kinase–dependent chromatin segregase^26^ and as a co-operative factor in the loading of the histone H3 variant CENP-A into centromeric nucleosomes^27^.

Regulation of the genome by bromodomains in kinetoplastids is of special interest due to the unusual polycistronic arrangement of the genome^28^ . As a limited number of transcriptional start regions controls large blocks of functionally unrelated genes, disrupting bromodomain function can have potent effects. For example, inducible deletion of BDF5 in *Leishmania mexicana* leads to a global reduction of transcription by RNA polymerase II, and the chemically tractable nature of its tandem bromodomains makes BDF5 a potential target for novel anti-leishmanials ^29,30^.

Given this context, we sought to identify ATAD2-like proteins in *Leishmania mexicana* and investigate their function. We identified bromodomain factor 7 as an ATAD2 orthologue and found that it is essential for promastigote to amastigote differentiation.

While it appears to have a weak association with chromatin, gene expression was altered in a Δ*bdf7* mutant leading to disruption of the stringent expression of the metabolic factors required for amastigote survival.

## Results and Discussion

### BDF7 is an orthologue of ATAD2

As part of a previous study of bromodomains in *Leishmania* species a custom HMM (Hidden Markov Model) was used to find divergent parasite bromodomains with weak sequence homology to other eukaryotes^31^. This revealed a previously unidentified bromodomain factor, that also contained an ATPase domain and an AAA+ lid domain – it was termed BDF7 for consistency with previous nomenclature (**Fig. 1A**). The arrangement of the domains suggested BDF7 is homologous to human ATAD2. The OrthoMCL DB resource places both *BDF7* and *ATAD2* in the same orthogroup (OG6_102094) making it likely that these genes share a common ancestor^32^. In *Leishmania mexicana* the gene ID is Lmx.11.0910, which encodes a protein of 1549 amino acid residues. Syntenic orthologues are found in *Leishmania donovani*, *Trypanosoma brucei* and *Trypanosoma cruzi*. A recent study provided ChIP-seq and immunoprecipitation datasets for *T. brucei* BDF7^33^, however, this study did not include a functional assessment of the protein.

**Figure 1:**
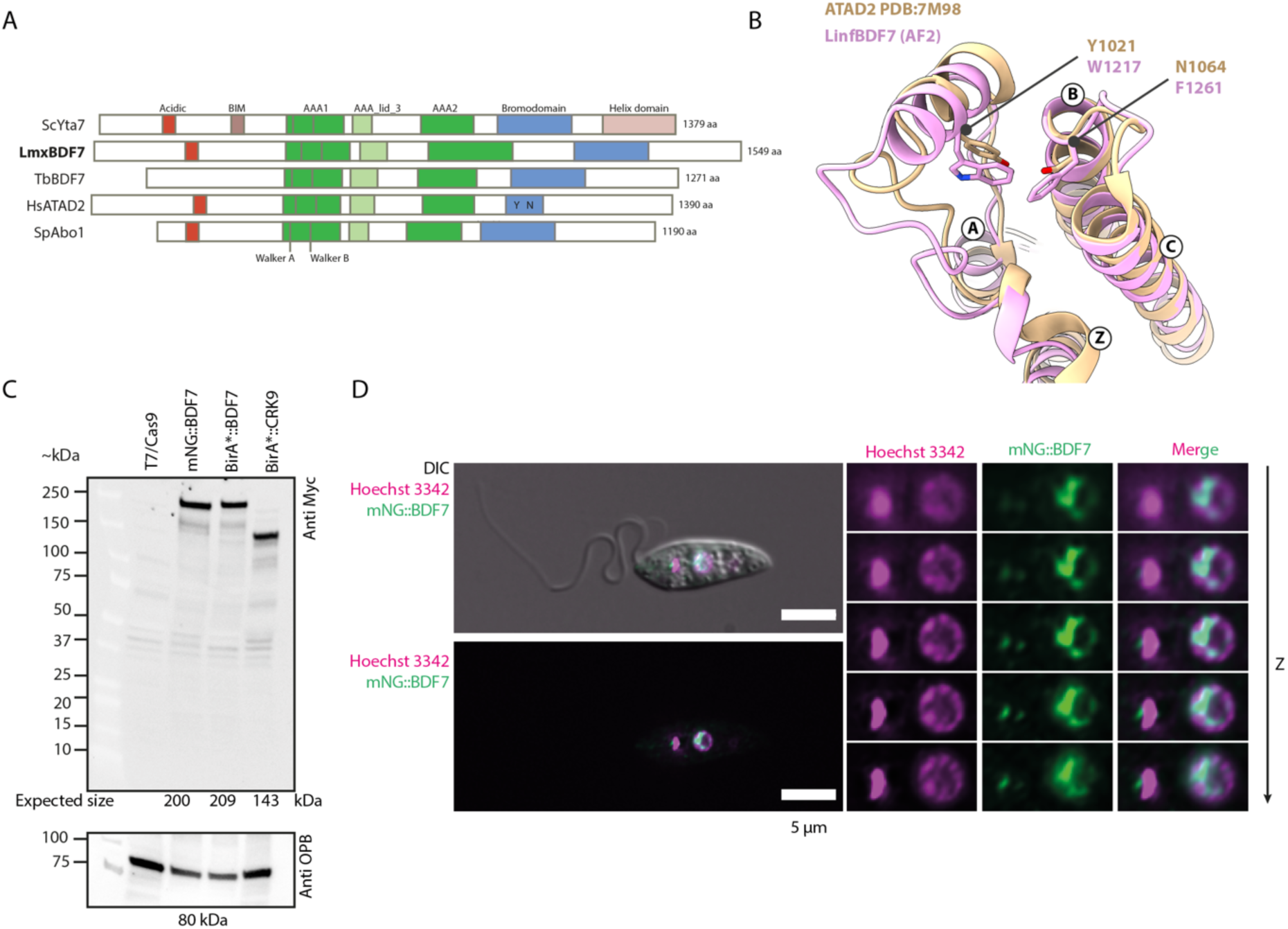
BDF7 protein architecture, localisation and expression. A. Overview of ATAD2-like protein architectures guided by sequence alignments to centre the ATPase domain. B. Superposition of X-ray crystal structure of HsATAD2 bromodomain (Tan- PDB: 7M98) with model of *L. infantum* BDF7 (Pink- AF-A4HV00-F1-v4, AlphaFoldDB) focusing on the acetyl-lysine binding pocket. In this instance *L. infantum* BDF7 was used due to it being available in AlphaFoldDB. Helices are labelled Z, A, B and C according to bromodomain convention. The conserved tyrosine and asparagine residues that mediate binding of acetylated lysine side chains are labelled in Tan for ATAD2. The corresponding residues of *L. infantum* BDF7 are labelled in Pink- the same residues are present in the *L. mexicana* and *T. brucei* sequences. The *L. mexicana* BDF7 bromodomain is 94.7% identical to L. infantum bromodomain with 98.2% similarity. C. Western blot to detect the 3xMyc epitope in BDF7 and CRK9 lines endogenous tagged with mNeonGreen (mNG) or BirA*. OPB in the lower panel is used as a loading control. Expected band sizes in kDa are noted under the individual blots. D. Live-cell, widefield fluorescence microscopy of *Lmx T7/Cas9 mNG::BDF7.* Promastigote cells were immobilised in CyGel and imaged in multiple z-planes allowing deconvolution analysis. The left panel represents the central slice showing the whole cell, with the right-hand panels showing a series of Z-planes.

In the absence of an experimental structure of the BDF7 bromodomain^31^ we used AlphaFold modelling to assess its predicted conformation which reinforced its atypical nature, it is evident that the conserved Y and N residues are replaced by tryptophan 1218 and phenylalanine 1262 respectively in *Leishmania* BDF7 (**Fig. 1B and Supplemental Fig. 1).** These bulky amino acid side chains project into the acetyl-lysine binding pocket such that acetylated peptides would be unable to bind in a conventional manner. Bulky residues are conserved at these positions in other kinetoplastids. The bromodomain of Yta7 has an extensive insertion in the ZA loop of the bromodomain and lacks the Y and N residues, again rendering it a non-canonical bromodomain^14^ . Thus, there is precedent for ATAD2 like proteins with non-canonical bromodomains.

ATAD2 has two AAA ATPase domains (AAA1 and AAA2) with an intervening AAA-lid domain (**Fig. 1A**). Using PFAM or INTERPRO searches, the AAA1 domain was readily identifiable in BDF7 although the second appeared to be more difficult to reliably find and is likely inactive. Alignments of the sequence of the AAA1 domain of BDF7 with sequences of other ATAD2 homologues showed a high level of conservation (59% identity and 76% similarity to ATAD2 for the residues used in Supplemental Fig. 2). The Walker A (GPPGTGKT) and Walker B (DEVDG) motifs were readily identifiable, indicating that this could be an active ATPase (**Supplemental Fig. 2**). Interestingly, there is a 14 amino acid residue insertion after the Walker A motif, which is absent in the other sequences including that of *T. brucei* BDF7. The Walker motifs were not found in the second AAA domain, and it is likely that AAA2 is inactive as an ATPase, as is the case for other ATAD2 like proteins^12^. AlphaFold3 modelling of the first AAA domain, the AAA lid domain and the second AAA-like domain suggested that the most likely oligomeric assembly of BDF7 would be a hexamer– consistent with the mammalian and yeast orthologs (**Supplemental Fig. 3A**).

Searching the LmxBDF7 sequence for the kinetoplastid nuclear localisation sequence^34^ (K[K/R]x[K/R]) identified 2 motifs that could potentially function as an NLS signal (residues KRGR^9–12^ & KKSR^200-203^) so we therefore sought to localise the protein by endogenous tagging and microscopy.

### Localisation of endogenously tagged mNeonGreen::BDF7

To test the hypothesis that BDF7 is a nuclear protein, we sought to localise it using the well-established *Leishmania mexicana T7/Cas9* strain (*LmxT7/Cas9*)^35^ and endogenously tagging BDF7 at its N-terminus with an mNeonGreen tag. Thus we generated a *3xMyc::mNeonGreen::BDF7* strain (*mNG::BDF7*) using reagents listed in Supplemental Table 1. To confirm the desired tagging of the target protein, cell lysates from mid-log promastigote cultures of the *mNG::BDF7* strain and the parental strain were resolved by SDS-PAGE and probed with an anti-myc antibody in a western blot (**Fig. 1C**). This revealed a single dominant protein which was consistent with the ∼200 kDa predicted molecular weight of the mNG::BDF7 fusion protein. No myc signal was detected in the lysate from the LmxT7/Cas9 parental strain in the 200 kDa range. Mid-log stage promastigotes of the *mNG::BDF7* strain were imaged after immobilisation in CyGel, using live-cell, widefield epifluorescence microscopy. Z-stacks of images were acquired to allow for deconvolution analysis to increase resolution (**Fig. 1D**). This revealed mNG signal predominantly, but not exclusively, in the nucleus of the cell. The signal was broadly distributed in the nucleus, and relatively heterogeneous, sometimes forming a ring around the nuclear periphery, or around and in structures that would likely be the nucleolus (**Fig. 1D, Supplemental Fig. 4**)^36^. Most cells also showed several small puncta, or a bar, of mNG signal close to kinetoplast DNA. Cells undergoing division were observed to have mNG signal in structures that could represent the mitotic spindle (**Supplemental Fig. 4B**).

Phosphorylated forms of BDF7 have been previously reported to be in proximity to the Mitotic Spindle Kinase (MSK) with several phosphosites were identified in the N-terminal region of BDF7 corresponding to residues S51, S75 and S126^37^ . Notably, in the Yta7 protein of *S. cerevisiae* phosphosites in the N-terminal region is reported to increase the ATPase activity and are important for progression through S-phase of the cell cycle^23,26^.

High-resolution immunofluorescence microscopy conducted on fixed promastigotes expressing a version of endogenously tagged 3xHA::BDF7 showed co-localisation of some of the BDF7 signal with the mitotic spindle **Supplemental Fig. 4C**. This confirms the observation of BDF7 at the mitotic spindle and opens the possibility that BDF7 function could be regulated by protein kinases.

BDF7 is not essential for promastigote viability but is essential for amastigotes to infect murine bone-marrow derived macrophages

BDF7-null mutants (Δ*bdf7*) were previously generated and validated as part of a Cas9-facilitated knockout screen to identify essential bromodomains^29^, and two clones were selected here for growth analysis. A very slight delay in proliferation was found in both clones when compared to the parental strain but all the cultures reached similar densities (∼10^7^ cells per ml) after 7 days in culture (**Fig. 2A**). Day 10 stationary phase cultures were next used to infect murine bone marrow derived macrophages to assess their capacity to survive and proliferate as amastigotes within a host cell. The *LmxT7/Cas9* strain was able to successfully infect macrophages (**Fig. 2B, 2C**), whilst both Δ*bdf7* clones exhibited a rapid decline in the percentage of macrophages that were infected by one or more parasites. By 72 hours only ∼5% of macrophages were infected, with this dropping further by 144 h post - infection (Fig. 2B, 2C). When the number of amastigotes per macrophage was counted this showed that the *Δbdf7* mutants infected macrophages with the same efficiency. In the subsequent timepoints the Δ*bdf7* strain failed to proliferate even in cells where amastigotes were visible (**Fig. 2D**). At 144 h no infected macrophages were observed for the Δ*bdf7* clone 1 and very few were infected by clone 2. This indicates that the strain is severely compromised in its ability to reach or survive as amastigote forms and for the successful infection of macrophages. To explore this further we used an Alamar blue cell viability assay to profile the viability of axenic amastigotes in Schneider’s media over six days. The *Δbdf7* clone 1 strain suffered a rapid and significant loss of viability compared to the *T7/Cas9* parental control (**Fig. 2E**). This suggests that BDF7 is important for an essential element of the amastigote’s intrinsic biology and not something extrinsic relating to host/pathogen interactions during macrophage infection. A derivative of Δ*bdf7* clone 1 was generated in which BDF7 was re-expressed from a linearised pRIB vector that was stably integrated into the RIB locus (**Supplemental Fig. 5)** Overexpression of the *BDF7* mRNA was confirmed by RNA-seq (see below), showing ∼60-fold expression of the mRNA. During *in vitro* amastigote differentiation conditions this BDF7 addback strain had the same viability as the parental T7/Cas9 strain (**Fig. 2E**), indicating that the phenotype in the Δ*bdf7* strain was specific to the lack of the *BDF7* gene and not from an off-target effect of the Cas9-assisted knockout process.

**Figure 2:**
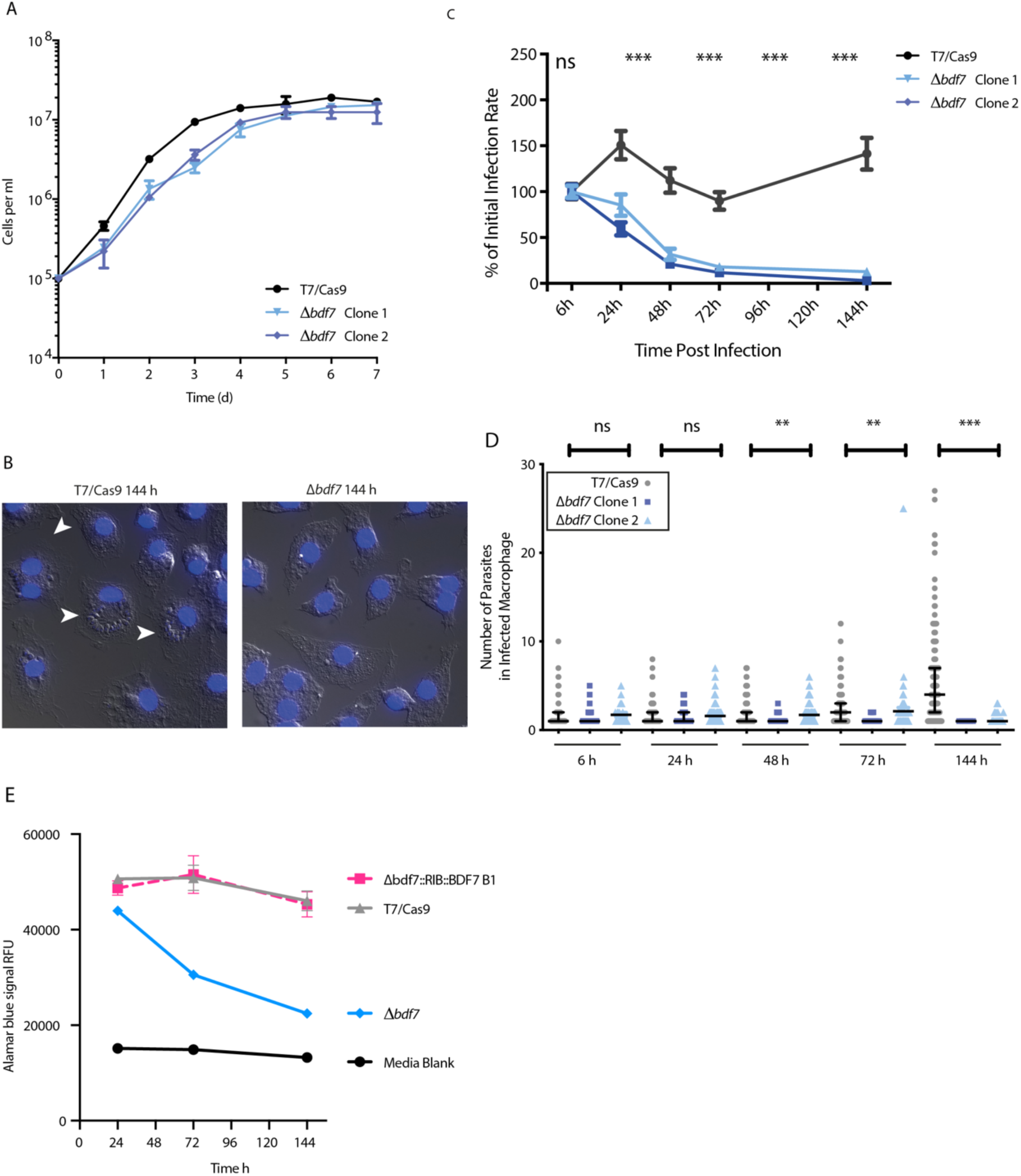
Characterisation of the BDF7 null mutant. A. Growth curve of promastigote cultures in HOMEM at 25 °C for Lmx T7/Cas9 and two independent Δ*bdf7* clones. Triplicate samples were counted for the parental control, duplicate samples were counted for each of the *Δbdf7* clones. Points show the mean and error bars show the range of the values. B. Example micrographs of macrophages containing intracellular Lmx T7/Cas9 amastigotes (left panel) compared to an image of macrophages infected with Δ*bdf7* parasites (right panel), after 144 h incubation. Imaging conducted using Zeiss AxioScope equipped with x63 DIC objective, imaging in transmitted white light and epifluorescence in the DAPI channel. Infected macrophages are denoted by white arrowheads, DAPI staining of DNA is denoted as blue pseudo-colour. C. Relative infection rate of mouse bone marrow-derived macrophages with *L. mexicana* T7/Cas9 or Δ*bdf7* line. Percentage of macrophages containing intracellular *Leishmania* over a 6-day infection time-course, the initial infection rate at T=6h was normalised to 100% for each cell line. Data points denote mean and standard deviation, these were compared by repeated T-test with Holm-Sidak multiple comparison correction, significant difference is indicated as (*=p<0.05, **=p<0.005 ***=p<0.0005), N=3, 100 macrophages counted per experimental timepoint. D. The number of intracellular *Leishmania* in each infected macrophage. Individual data points are denoted, with bars for median and interquartile range, data were analysed by Kruskal-Wallis test with multiple comparison corrected by Dunn’s method, (*=p<0.05, **=p<0.005 ***=p<0.0005), n=100 macrophages counted, N=3 technical replicates (wells). E. Alamar blue cell viability assay for axenic amastigote differentiation cultures. Points represent the mean of six replicate wells, bars represent the standard deviation.

### ChIP-seq suggests low occupancy of chromatin by BDF7

As BDF7 is predicted to have ATAD2-like properties as a chromatin regulator/remodeller, we adopted a ChIP-seq approach to identify which regions and functional elements of the genome it could interact with. ChIP profiles of ATAD2-like proteins in different experimental systems show considerable heterogeneity. ATAD2 has a canonical bromodomain and when ChIP-seq was performed on embryonic stem cells it showed enrichment over gene-containing regions of the genome (exons and introns) while intergenic regions lacked ATAD2 enrichment. ATAD2 bound chromatin with high-affinity and could even be precipitated under non-crosslinking conditions^15^ . Yta7 was shown to bind chromatin, in ChIP-chip assays, localising mainly to intergenic regions^22^. Notably, Yta7 had chromatin binding activity even when the bromodomain was deleted entirely^22^. Indeed, BDF7 from *T. brucei*, which lacks the conserved acetyllysine binding residues, **Supplemental Figure 8C**, has been shown to bind to the spliced-leader RNA locus and a subset of genomic loci where pol II transcriptional termination regions contain pol III expressed non-coding RNA genes^33^.

We generated a new strain (*Lmx 10xTY::BDF7*) with BDF7 tagged by an N-terminal 10xTY epitope (EVHTNQDPLD) so that it lacked lysine residues^35,38^ . Western blot analysis of the resultant cell line showed a prominent protein at a size consistent with BDF7 (∼200 kDa). Despite the presence of a weaker band (ca 37 kDa) cross-reactive with the parental *LmxT7/Cas9* strain, we pursued ChIP-seq analysis with these cell lines (**Fig. 3A**). ChIP was performed using an anti-TY antibody and protein G magnetic beads on formaldehyde crosslinked chromatin from both the 10xTY::BDF7 strain and the parental control strain.

**Figure 3:**
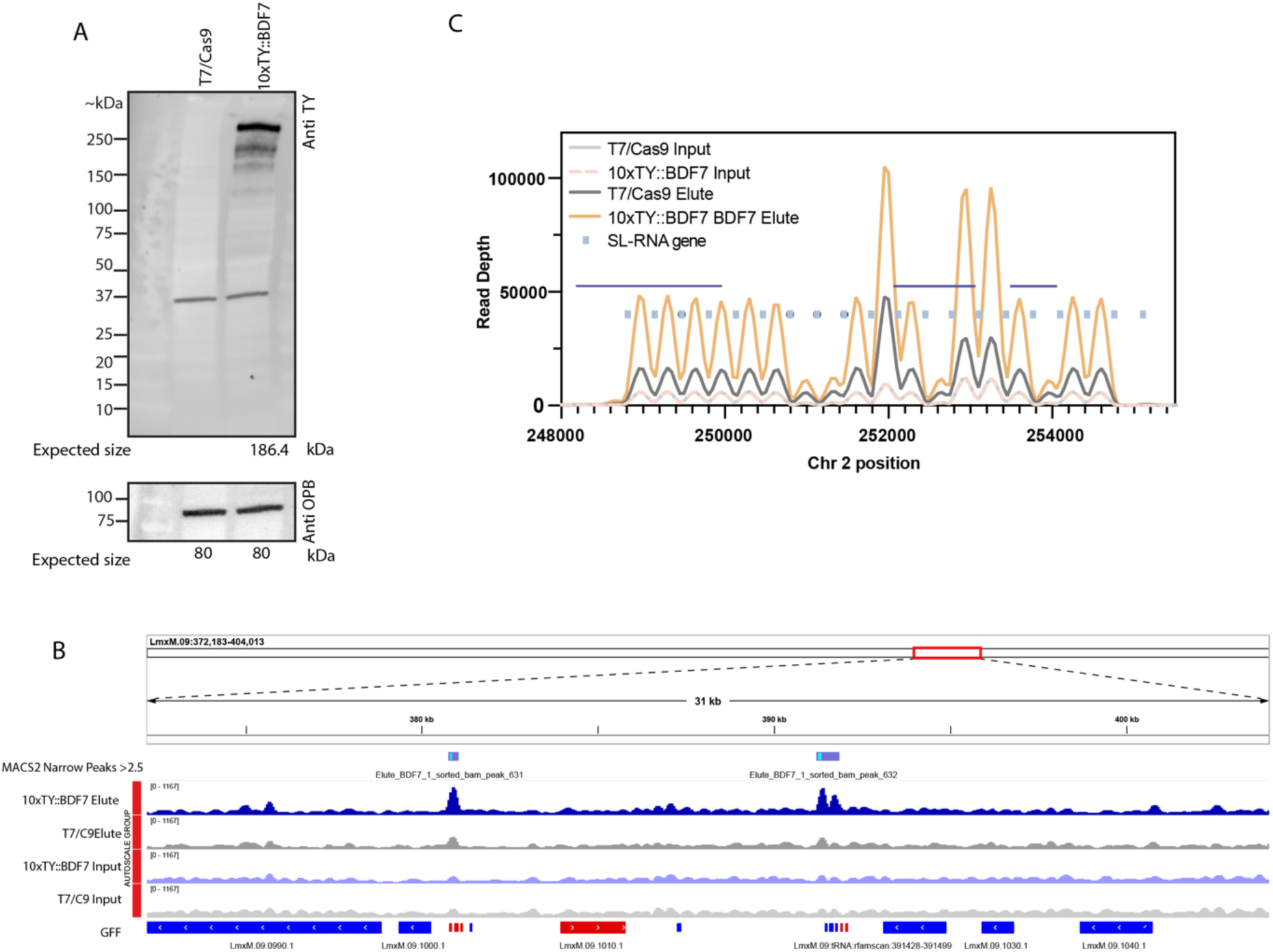
Generation of 10xTY::mNG strain and ChIPseq. A. Western blot with anti-TY antibody to detect 10xTY::mNGBDF7 generated by endogenous tagging. The lower panel uses OPB as a loading control. The expected size for 10xTY::BDF7 is 186.4 kDa , for OPB it is 80 kDa. B. Genome browser (IGV) view of a portion of Lmx Chr9, showing a divergent and convergent strand switch region, each containing a cluster of rRNA and tRNA genes. The average depth of sample sequencing reads are shown for the 10xTY::BDF7 ChIP and control ChIP, and the respective inputs are shown, these are the averages of 3 replicates for each sample condition. The top track shows the MACS2 output, highlighting regions that are statistically enriched for BDF7 but not crossing the 5-fold cut-off to meet the MACS2 “Robust” threshold. C. Average read depth from triplicate ChIP input and elution samples from anti-TY pulldown on LmxT7/Cas9 and 10xTY::mNG strains mapped against LmxT7/Cas9 Nanopore assembly, focused on the spliced leader gene array on Chromosome 2. The positions of individual spliced leader genes are indicated using blue rectangle, the regions called as enriched by MACS2 are denoted by the purple horizontal lines.

Input and elution samples were submitted for Illumina sequencing and reads were initially aligned to the *L. mexicana* U1103 reference genome^39^; the BAM alignment files were then used in a MACS2 ^40,41^ analysis to search for areas of the genome enriched in the BDF7 pulldown. The Elution samples from the *LmxT7/Cas9* strain provided the control samples to establish the λ local values (local background). Using the default lower limit of fold enrichment in MACS2 (5-fold), no regions met the threshold of being enriched for BDF7 (**Supplemental Table 2**). Numerous regions were found to meet statistically significant levels of enrichment that were under the 5-fold cut-off, but given the low-fold enrichment the biological significance of these is unclear. Several peaks were identified in strand switch regions with rRNA and tRNA genes, suggesting a similar distribution to TbBDF7 but this was not consistent across all of them (**Fig. 3B, Supplemental Fig. 6**) Comparison of the data with an alternative method, deepTools bamCompare with SES (Signal Extraction Scaling) normalisation also failed to highlight any consistent areas of enrichment^42^ (**Supplemental Fig. 6**).

When BDF7 was assessed by ChIP-Seq in *T. brucei* bloodstream forms using a mCherry::BDF7 tagged allele, it was found to be enriched at a subset of 20 of the 154 transcriptional termination regions (TTR), specifically those where the TTR coincided with an RNA pol III transcribed gene – e.g. tRNA or snoRNA genes and also found enriched at the spliced-leader RNA locus (SL-RNA array)^33^ so we assessed this in our assay. As the *L. mexicana* U1103 reference genome is a short-read assembly the SL-RNA is collapsed, we sought to assess potential enrichment at the SL-RNA array. The reads were therefore mapped against a genome assembly derived from long-read Nanopore sequencing^43^.

Viewing the reads mapped against this region showed there was an increase in the amount against the SL-RNA intergenic regions, which MACS2 called as significantly enriched albeit with high background and low signal - the enrichment was only in the region of 1.4-1.5-fold (**Fig. 3C**). Notably the first peak in this region occurred after the first SL-RNA gene, which could be consistent with the observation that nucleosomes are found after the SL-RNA and that the gene body and promoter are devoid of nucleosomes, to which it would be predicted that BDF7 binds^44^.

Histones, histone modifications and variants^45^, and chromatin bound factors^46^ in *Leishmania* have been amenable to ChIP-seq, including bromodomain factor 5^29^. We utilised a similar protocol as for BDF5-ChIP suggesting that the challenges to identify BDF7-enriched loci was not a general failure of the method and reflects a state where BDF7 is not tightly engaged with chromatin in *Leishmania*. The low ChIP signal is likely reflective of the lack of canonical bromodomains in BDF7 and, given a potential role of the protein in nucleosome dissociation, this would lead to it have only transient interactions with chromatin. Considering this, we next characterised BDF’s local protein landscape with proximity-biotinylation proteomics.

### XL-BioID analysis of BDF7 proximal proteins

Our lab has previously developed cross-link proximity biotinylation (XL-BioID), to identify components of chromatin associated complexes, such as the BDF5-containng CRKT complex and the kinetochore, in *Leishmania* ^4748^. We reasoned that XL-BioID would be an appropriate technique to apply to BDF7 given its discrete nuclear location (**Supplemental Fig. 7**). BDF7 was tagged at its N-Terminus with 3xMyc::BirA* tag, giving the cell line *BirA*::BDF7*. To subtract endogenously biotinylated proteins in the XL-BioID data analysis, and non-specifically labelled proteins we generated an organellar control strain expressing the nuclear localised protein kinase CRK9 (BirA*::CRK9), an essential protein kinase that regulates mRNA splicing^29,49,50^. Both strains also encode a 3xMyc epitope tag prior to the BirA* fusion protein, allowing for ease of detection using anti-myc antibodies. Western blot analysis of each cell line (**Fig. 1B and Fig. 4A**) showed a single dominant band, migrating at the expected molecular weight (209 kDa and 143 kDa for BirA*::BDF7 and BirA*::CRK9 respectively). These were then grown in cultures with or without additional D-biotin supplementation (150 µM) overnight and the biotinylation signal identified in a western blot (**Fig. 4A**). Fluorescence signal identified bands in both strains consistent with the natively biotinylated proteins found in *L. mexicana.* Additionally, prominent bands consistent with BirA*::BDF7 and BirA*::CRK9 were observed in the respective strains. These bands overlapped with anti-myc, indicating successful biotinylation of the bait proteins.

**Figure 4.**
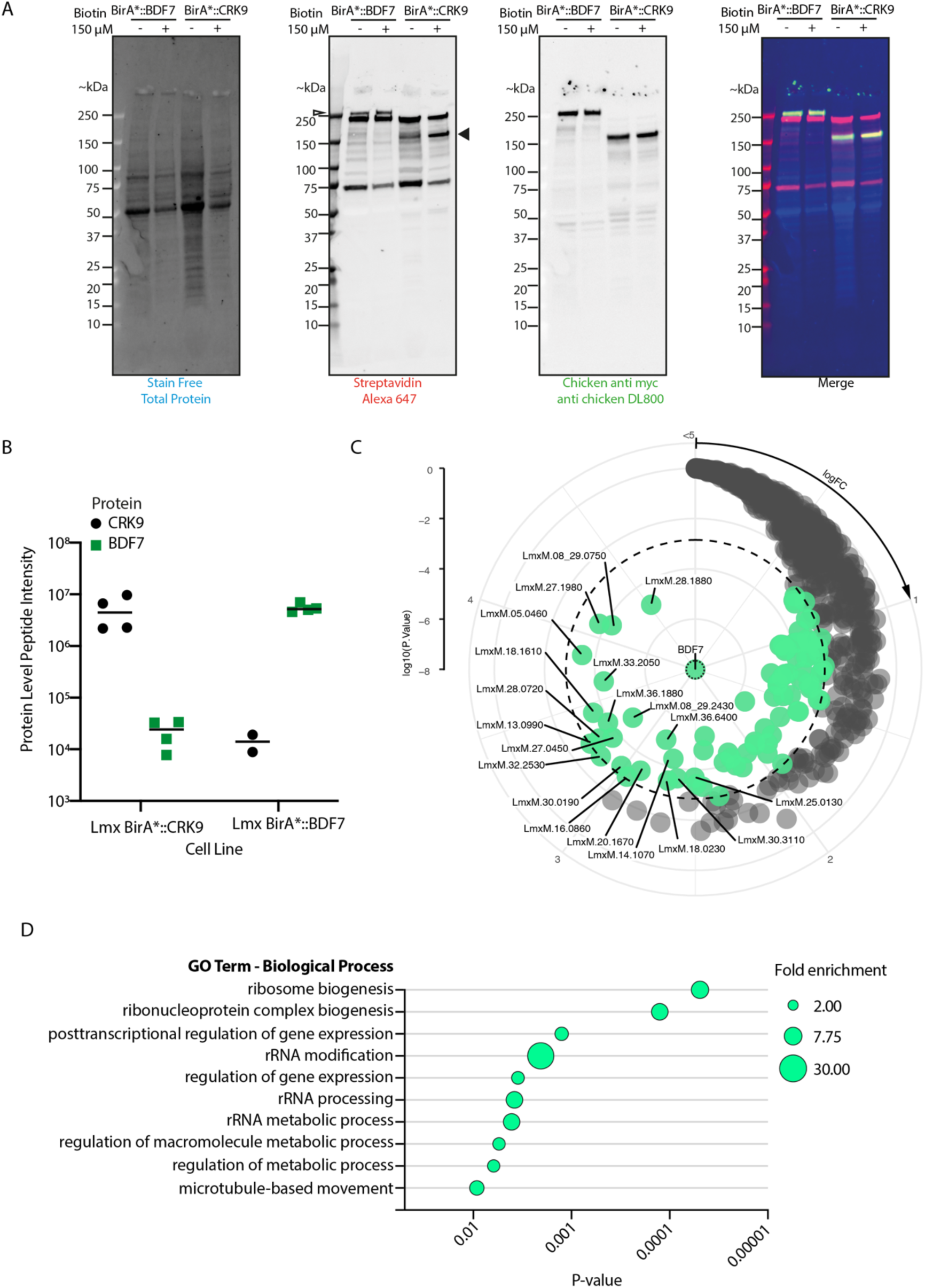
*XL-BioID to reveal the BDF7 proximal proteome.* A. Western blots and streptavidin blots showing the BirA*::BDF7 and BirA*::CRK9 strains response to overnight treatment with 150 µM biotin. BioRad Stain Free total protein labelling is used as a loading control. In the streptavidin panel a narrow arrowhead on the left edge indicates the expected size of BirA*::BDF7 (209 kDa), on the right-hand edge a broad arrowhead indicates the expected size for BirA*::CRK9 (143 kDa). The other bands represent other biotinylated proteins in the sample (endogenous or proximal to the bait protein). The Chicken anti-myc panel shows directly the tagged bait proteins. All three channels are then merged in the right hand panel. B. Chart showing the protein level peptide intensity of CRK9 and BDF7 in the respective sample groups. Individual points indicate the value for one of each quadruplicate sample, and the line indicates the mean. There were 2 replicates with missing values for CRK9 in the BirA*::BDF7 therefore only 2 points are plotted. C. Radial plot depicting BDF7 proximal proteins. Green points represent significantly enriched proteins (> 2-fold , p-adj<0.01), grey are detected proteins that are not statistically significantly enriched in proximity to BDF7. Dotted ring indicates the p-value cutoff. Clockwise on the circumferential axis indicates increasing fold-change, the radial axis indicates the log10(P Value). Selected hits (over 5.5-fold enriched) are annotated with gene ID. D. The top 10 GO Terms (Biological Process) associated with the 86 significantly enriched proteins are plotted to depict the P-value on the X-axis, with the size of the circle reflecting the fold enrichment of those terms. GO term enrichment analysis was conducted with TriTrypDB Analyse Results function using the gene ID list as an input.

Biotinylation of the bait proteins was observed in cultures without the addition of extra biotin although in the BirA*::CRK9 strain this signal did appear to be more responsive to the addition of extra biotin. While this lack of inducibility may hinder dynamic, pulse/chase-type experiments we deemed it acceptable for a cataloguing approach for initial characterisation of BDF7 proximal proteins.

A full-scale XL-BioID workflow was conducted using four replicates of each cell line and the streptavidin enriched proteins were on-bead digested using trypsin/Lys C prior to desalting and mass spectrometric analysis. In this instance, we used the mass spectrometer in label-free data independent acquisition mode to quantify peptide intensities, allowing for greater proteome coverage, faster run times and reduced missing values^51^. Initial assessment of the signal coming from the target proteins in each separate pulldown showed good separation of BDF7 and CRK9 signal, indeed signal from CRK9 was only detected in 2 of the BirA*::BDF7 samples confirming that the two proteins are spatially separate and not co-enriched (**Fig 4B**). These data were processed using missing value imputation, then analysed with limma^52^ to identify differentially enriched proteins in each sample set, revealing proteins found in proximity to each bait protein. Overall, the analysis identified 1554 proteins between the two sample groups, of which 86 were determined to be significantly enriched in proximity to BDF7 and 131 in proximity to CRK9, using cutoffs of log2FC>0 and 1% FDR (**Fig. 4C**, **Supplemental Table 2**). Proximal interactions were successfully identified by the workflow, exemplified for CRK9 by the interacting partner CYC12 being the second most enriched protein after the CRK9 bait (**Supplemental Fig. 7**)^50,53^.

BDF7 was the most enriched protein in the BirA*::BDF7 sample group, being enriched log2FC 8.14. The 86 BDF7 proximal proteins were assessed using GO term analysis, the top 10 biological process terms associated with these proteins were linked to processes such as ribosome and ribonucleoprotein complex, post-transcriptional regulation of gene expression (**Fig. 4D**). Eight proteins were detected as being >10-fold enriched, BDF7, LmxM.28.1880 - Nucleolar protein 136, putative, LmxM.08_29.0750 - U3 small nucleolar ribonucleoprotein protein MPP10, putative, LmxM.27.1980 - FtsJ cell division protein, putative, LmxM.05.0460 - GTPase, putative, LmxM.33.2050 - ATP-dependent RNA helicase, putative, LmxM.18.1610 - Noc2p family, putative, LmxM.36.1880 - CBF/Mak21 family, putative. NOP136 has been identified in *T. brucei* as a nucleolar protein associated with the mitotic spindle, it has homology to BMS1, an ATPase in humans and brewer’s yeast that is a component of the ribosome small subunit (SSU) processome^54^ . MPP10 is also a component of this complex^55^ . LmxM.27.1980 is annotated as FtsJ-like protein, which are methyltransferases that can methylate RNA, including ribosomal RNA. LmxM.05.0460 has homology to Nucleolar GTP-binding protein 2 a GTPase that associates with pre-60S ribosomal subunits^56^ . LmxM.33.2050 may be a DEAD-box RNA helicase, with some homology to DBP10 of yeast, which again is involved in processing 60S ribosomal subunits. LmxM.18.1610 - Noc2p ^57^ . In *S. cerevisiae* this has been shown to associate with Mak21, a homolog of which (LmxM.36.1880) was also identified in the XL-BioID analysis. These proteins are typically found in the nucleolus, for example NOP136, MPP10, FtsJ, Noc2p and the orthologs of LmxM.05.0460, LmxM.05.0460 are identified in the TrypTag dataset as being nucleolar; and Mak21 is found in the nucleolus in LeishTAG imaging.

The remaining list of proximal proteins contained a range of factors involved in different processes such as RNA binding proteins, poly(A)-binding proteins, kinesins and cytoskeletal proteins, as well as some metabolic enzymes (**Supplemental Table 2**). ATAD2 has been purified from mouse embryonic stem cells using tandem-affinity purification and ChIP, and the interacting proteins assessed by mass spectrometry^15^. This indicated that ATAD2 pulldown would also retrieve histones, the FACT complex, and other proteins involved in chromatin remodelling, DNA repair and replication, splicing, cohesion and kinesins. In combination with the broad spectrum of proteins that were retrieved in the ChIP proteomics and the fact that ATAD2 had never been identified in an essentiality screen the authors concluded the ATAD2 plays a generalist “helper” function, such that its loss can be compensated by more specialised factor. Our results, while retrieving different classes of proteins, indicate that the BDF7 proximal environment also includes a diverse set of proteins.

A previous study of chromatin binding factors in *Trypanosoma brucei* conducted immunoprecipitation and mass-spectrometry of mCherry tagged BDF7 in *T. brucei* bloodstream forms and reported interactions with only three proteins – nucleosome assembly proteins 1-3. The *L. mexicana* NAP orthologues (LmxM.08_29.2340, LmxM.19.0440, LmxM.20.1290, and LmxM.30.1750) were detected in the DIA dataset but were not identified as differentially enriched between the two sample groups. This could indicate that BDF7 is not specifically associated with NAPs in *Leishmania* or that the CRK9 spatial control is also in proximity to the NAPs. Additionally, while the TbBDF7/NAP association might be quite stable, the IP approach used by Staneva et al. would not exclude post-lysis interactions. Histones were detected in our analysis, but none were specifically enriched with BDF7. An immunoprecipitation study of *A. thaliana* BRAT1 did also not identify NAPs, and showed BRAT1 interacted with another ATPase BRP1 which contains a plant homeodomain-like zinc finger and an AAA-type ATPase^58^, no proteins homologous to this were found in the BDF7 proximal proteome.

Intriguingly, the enrichment of components important for ribosome biogenesis may reflect the nucleolar/peri-nucleolar localisation of BDF7 observed in *Leishmania* (**Supplemental Fig. 4**). The importance of stage specific ribosomes and ribosome modification are starting to be revealed in kinetoplastids, with even single pseudouridine modifications proving impactful on stage-specific translation and parasite fitness^59,60^ .

Therefore, any effect of BDF7 deletion on ribosome biogenesis that may be tolerated in promastigotes could still have a deleterious effect in amastigotes.

### RNA-seq to evaluate the *Δbdf7* mutant during differentiation

Given that BDF7 was not essential for the growth and survival of promastigotes, but that the BDF7 deletion mutant failed to grow as amastigotes forms, we decided to conduct an mRNA-seq experiment to assess the cellular states of the mutant during this process and identify potential causes for transition failure. Cultures of the *LmxT7/Cas9*, the Δ*bdf7* and the Δ*bdf7::BDF7* strains were set up in Graces medium (pH 5.5), an acidic medium which promotes metacyclogenesis, and grown at 25 °C for 7 days ^61^. At day 7 the flasks were shifted to 35 °C for 24 h to provide the second signal to trigger amastigogenesis^62^. Samples were taken at day 3, day 7 and day 8 for RNA extraction and mRNA sequencing (**Fig. 5A**).

**Figure 5:**
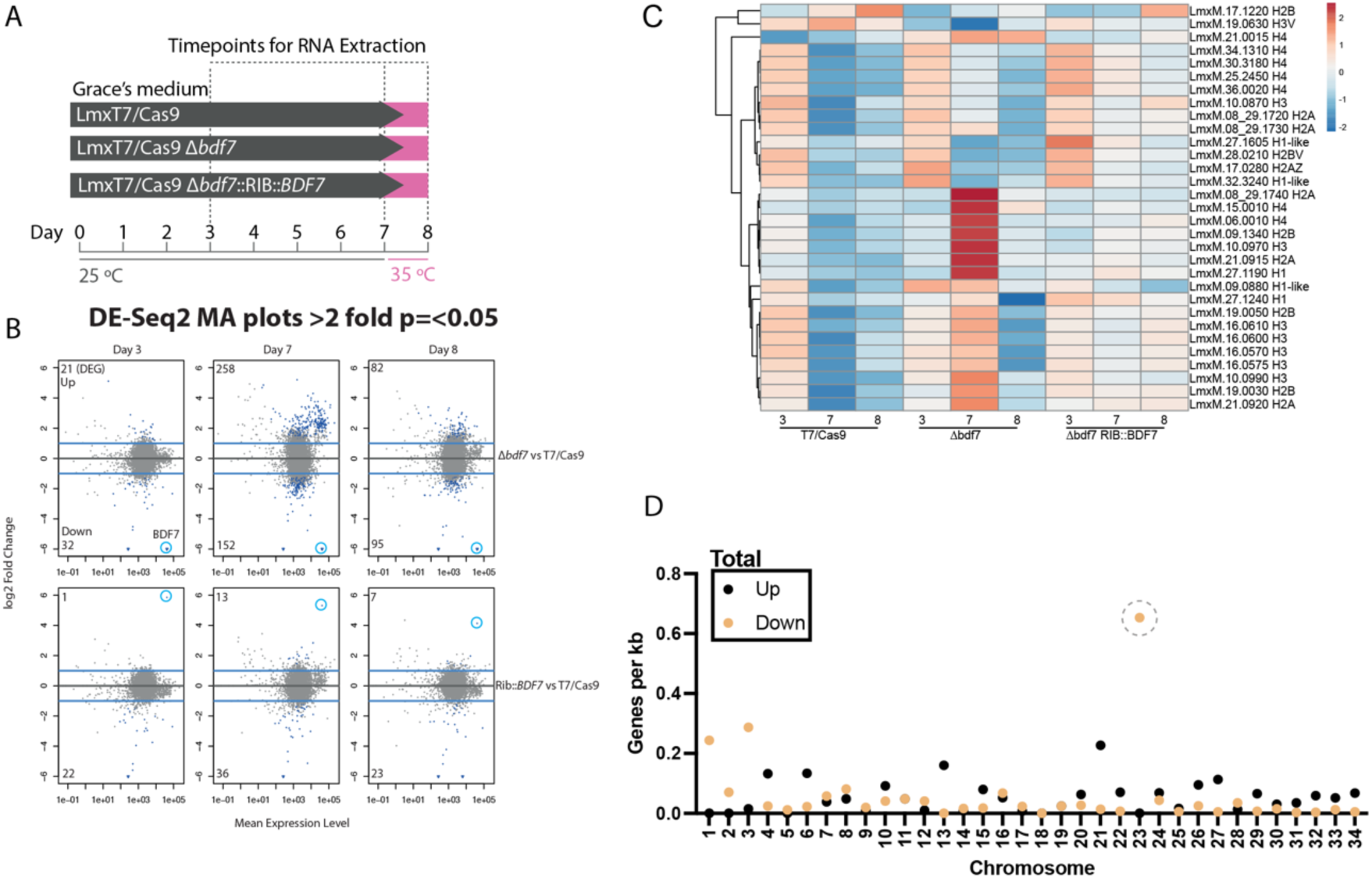
RNA-seq on BDF7 null mutants. A. Schematic showing the strains used, the media (Grace’s) and temperature regime, and the sampling points. B. MA-plots from the DE-SEQ2 analysis. Comparisons are made between the Δbdf7 strain and Lmx T7/Cas9 (top row of plots) and Δbdf7::RIB::BDF7 against Lmx T7/Cas9 (lower row of plots). mRNAs showing >2-fold change (y-axis) with p(equal or less) 0.05 (x-axis) are highlighted in blue, the BDF7 point is highlighted by a light blue ring. C. Clustered-heatmap of histone genes, coloured by the intensity of the z-value. D. Interleaved scatter plot depicting the number of differentially expressed genes per chromosome, normalised by chromosome length to give number of DE genes per kb. The point indicating the amount of DE genes per kb on Chr 23 is circled by a dashed line to highlight it.

DESeq2 was used to identify differentially expressed genes in the mutants (**Fig. 5B**)^63^. Comparing the *Δbdf7* strain to the *T7/Cas9* parental strain at day 3 revealed that there were few differentially expressed genes; 21 up and 32 down with cut-offs of >2-fold change, p<0.05 (**Supplemental Table 3**). BDF7 was identified as the most downregulated gene, consistent with the strain being a null mutant. Of the 21 upregulated genes the most upregulated (log2 fold change 5.11) was Histone H4, the next most upregulated (log2FC 2.89) was a predicted pseudogene Lmx.27.1740). Intriguingly, of the 32 downregulated genes, 14 were located on chromosome 23. No other positional enrichment was noticed for this cohort, with genes being spread evenly over 13 different chromosomes (**Supplemental Fig. 8**).

Comparing the *Δbdf7::RIB::BDF7* strain to the parental strain at day 3 identified BDF7 as ∼60-fold upregulated, the only gene with increased expression levels and reflective of the high transcript levels driven by pol I transcription of RIB integrated vectors. In the *Δbdf7::RIB::BDF7* strain at day 3, 22 genes were significantly decreased (**Supplemental Table 4**). Therefore, it appears that the overexpression of the *BDF7* mRNA does not induce a detrimental transcriptomic state, and this strain also differentiates normally to axenic amastigotes (**Fig. 5B**).

At day 7, the comparison of the *Δbdf7* strain to the *T7/Cas9* identified 258 upregulated mRNAs and 152 that were decreased, while the *Δbdf7::RIB::BDF7* only had 13 upregulated and 36 downregulated compared to the *T7/Cas9* – showing the differences were emerging in the *Δbdf7* mutant were reversed in the addback strain (**Supplemental Tables 5 and 6**). At day 8, following transfer to 35 °C, 82 genes were upregulated and 95 downregulated in *Δbdf7* compared to *T7/Cas9*, the *Δbdf7::RIB::BDF7* strain had 7 upregulated and 23 downregulated when compared to *T7/Cas9* (**Supplemental Tables 7 and 8**). Intriguingly, the biggest difference in the *Δbdf7* mutant was at day 7, before the transition to amastigote differentiation conditions. This might suggest that the Δ*bdf7* strain was not pre-adapted to making the metacyclic-amastigote differentiation step. Therefore, the differences in the transcriptome at day 7 were assessed further.

GO analysis of the up-regulated genes in the *Δbdf7* strain suggested that processes involving translation, peptide biosynthesis and other anabolic activities were upregulated in the top 10 GO terms, with the “gene expression” biological process term also being present (**Supplemental Figure 9A**). Terms enriched in the downregulated genes included those associated with ion membrane transport, proteolysis, adhesion and amino acid biosynthesis (**Supplemental Figure 9B**). It is notable that ribosomal genes (which trigger the ‘translation’ GO term) are upregulated given that the XL-BioID data implicate BDF7 as proximal to proteins involved in ribosome maturation, and again indicates that the cell may be compromised in the content or type of ribosomes it possesses.

It was also noted that in the *Δbdf7* a subset of histone genes appeared to be differentially expressed compared to the *T7/Cas9* (**Fig. 5C**) and these also showed a different pattern of expression during the time course – which is intriguing considering the previous reports of Yta7/Abo1 in regulating the stability of nucleosomes and the expression of histones^21,23^ . The *L. mexicana* histone genes identified did not appear to form clear pattern based on genome localisation or histone subtype, with the exception of a small cluster of H3 genes on chromosome 16.

Genes on Chromosome 23 were again consistently identified as enriched in the analysis, with 50 of the 152 downregulated genes being on that chromosome – which contains 210 protein coding genes (**Supplemental Fig. 8B**). This is a striking observation as we are not aware of this type of positional effect being noted in other *Leishmania* mutants. Given a putative role of BDF7 in chromatin remodelling it may suggest that a chromatin structure or architecture specific to chromosome 23 is being disrupted in the *Δbdf7* mutant. Overall, Chromosome 23 held a large proportion of the downregulated genes– around 35% of all the DE genes at all time points (Fig. 5D). Chr23 is disomic (as opposed to Chr30, for example) so we looked for other distinguishing features. One striking feature is a very high concentration of non-coding RNAs in the convergent strand switch region between positions 217,000 and 220,000 bp. This cluster contains 10 tRNA genes, 1 rRNA gene and 1 unspecified ncRNA. Although our ChIP-seq analysis did not indicate enrichment of BDF7 at this site in *L. mexicana,* the syntenic locus in *T. brucei* was enriched for BDF7^33^ (Supplemental Fig. 8C). This region represents the boundary of a pol II termination site, and includes 3 dense clusters of tRNA genes, over each of which BDF7 signal was enriched. Chr 8 in *T. brucei* represents a fusion event^64^, and this region retains synteny with *Leishmania* Chr 23, with conservation of protein coding genes in the pol II cistrons and the tRNA cluster between them. Although the chromatin features that might act as insulators are poorly understood in *Leishmania*, tRNA genes have been proposed to correlate with insulator elements in the *Trypanosoma cruzi* genome^65^. Yta7 has also been shown to play an important role in establishing barrier elements in *S. cerevisiae*, including at tRNA genes^22,66^. Therefore, BDF7 loss might disrupt putative barrier elements in *L. mexicana,* following which, pol III could disrupt proper regulation of pol II transcribed genes on the rest of chromosome 23; thus creating circumstances where there is a broader, non-specific downregulation of mRNA from that chromosome. While the ChIP-seq results suggest BDF7 does not tightly associate with chromatin, it is still capable of exerting an effect on locus-specific gene expression.

Among the other genes with reduced expression levels at Day 7 and 8 (**Supplemental Figure 10**) (under amastigote inducing conditions) are those encoding glutamine synthetase and glutamate dehydrogenase. Glutamine is a key amino group donor, required for essential pathways such as pyrimidine and amino sugar biosynthesis. *Leishmania* amastigotes are known to undergo a pronounced shift in central-carbon metabolism when compared to promastigote forms. Both growing and stationary phase promastigotes utilise a wasteful method of metabolism in which large amounts of glucose and amino acids are transported into the cell but then excreted up to 70% of this only partially metabolised. Amastigotes, on the other hand, have a stringent metabolic response^67^; characterised by dramatically reduced capacity to uptake glucose and amino acids, but which are then utilised much more efficiently. A critical element of amastigote metabolism is the dependency on *de novo* synthesis of glutamine from glutamate, which in turn is derived from α-ketoglutarate generated by the citric acid cycle. As scavenging of glutamate is reduced by ∼30-fold in amastigotes ^67^, this would make these parasite forms sensitive to perturbations in glutamine synthesis. Our DEseq2 analysis revealed that the mRNA for glutamine synthetase is significantly reduced at Day 7 (-1.3 log2-fold) and decreases to -2.3 log2-fold reduced at the day 8 timepoint - this is likely to be a fatal complication for the amastigote forms. While glutamine synthetase expression trended lower in the *Δbdf7::RIB::BDF7* strain it was not significantly lower in the DESeq2 analysis, which may provide enough BDF7 to complete the transition.

## Conclusion

BDF7, while not essential for promastigotes of *Leishmania*, is critical for the survival of amastigote forms. In the context of literature about the roles for ATAD2-like factors, we propose that BDF7 is a multifunctional protein whose dysfunction leads to multiple consequences. We observed BDF7 in multiple locations in the cell/nucleus, with ChIP-seq showing low levels of association with chromatin. Our XL-BioID shows proximity to ribosome maturation factors, consistent with the nucleolar localisation observed for BDF7. This might be due to the relative abundance of those factors in the cell or their propensity to be biotinylated, or a bias introduced by using CRK9 as a spatial control. Meanwhile, RNA-seq data identified the consequences of BDF7 deletion on gene expression, indicating BDF7 has functions associated with chromatin compartmentalisation – for example insulating pol III promoters at transcriptional stop regions. Therefore, our different assays seem to capture differing roles of BDF7. The conclusion that BDF7 has generalist functions is consistent with the emerging consensus from studies of orthologues in other organisms. BDF7 null mutant promastigote forms are viable, matching results in yeast where null mutants of Yta7 are similarly viable. The yeast mutants are however sensitive in broad range assays that expose the cells to a stress or look for synthetic lethality. Parallels can be drawn here with the exposure of *L. mexicana* to acidic conditions and temperature shift. While these are part of the natural life-cycle of the parasite, and therefore may not be classically stressful, the parasite has to make specific alterations to its biochemistry and morphology.

Substrate profiling of ATAD2-like proteins has focused on nucleosomes, but given the degenerate nature of the BDF7 bromodomains it may be possible that BDF7 is not restricted to such substrates. Indeed, while we do not see a strong association with chromatin or nucleosomes in the ChIP-seq and XL-BioID respectively, we do see a signature of this in our RNA-seq analysis. We interpret this as BDF being able to play multiple, generalist roles in the cell, which may fine tune cellular environments for other specific factors to better function. Given that ATAD2-like factors are rarely identified in essentiality screening it is interesting that we find it essential for a specific life-cycle transition point in *Leishmania*. As the amastigote forms of the parasite are dependent on a finely balanced stringent metabolic response upon differentiation, our hypothesis is that small alterations to this balance, particularly glutamine flux, are caused by BDF7 deletion and prove fatal to the cell.

While essential for amastigotes, BDF7 contains a non-canonical bromodomain and we did not establish any evidence for the essential role of this domain - therefore this domain would represent a challenging target for anti-leishmanial development. However, the apparent range of potential functions for BDF7 highlights the need to further investigate ATAD2-like proteins in diverse eukaryotes. Further exploration of BDF7’s cellular functions could be focused on chromatin accessibility^68^ , transcriptional activity^69^ , and trying to identify direct substrates in its apparent noncanonical role in ribosomal protein biogenesis.

## Data Availability

### Proteomics

The results of the mass spectrometry are available at MassIVE.

Pre-publication access can be obtained from MassIVE with the following link: ftp://MSV000097088@massive.ucsd.edu

### Sequencing

RNA-seq and ChIP-seq reads are available at SRA under the BioProject ID PRJNA1330792.

## Author Contributions

**Conceptualization:** Nathaniel Jones, Félix Calderón, Jeremy Mottram,

**Formal Analysis:** Nathaniel Jones, Adam Dowle,

**Funding Acquisition:** Félix Calderón, Nathaniel Jones, Tony Wilkinson, Jeremy Mottram

**Investigation:** Nathaniel Jones, Alex Johnson,

**Methodology:** Nathaniel Jones

**Project Administration:** Raquel Gabarró

**Resources:** Adam Dowle

**Supervision:** Nathaniel Jones, Félix Calderón, Tony Wilkinson, Jeremy Mottram

**Visualization:** Nathaniel Jones

**Writing – Original Draft Preparation:** Nathaniel Jones

**Writing – Review & Editing:** All the authors

## Competing Interest Statement

Félix Calderón & Raquel Gabarró are employees of GlaxoSmithKline. This work was supported by funding from GSK through the Pipeline Futures Group, and a Fellowship from a Research Council United Kingdom Global Challenges Research Fund under grant agreement A Global Network for Neglected Tropical Diseases grant number MR/P027989/1. to Nathaniel Jones. This work was part-funded by the Wellcome Trust [ref: 204829] through the Centre for Future Health (CFH) at the University of York.

## Materials and Methods

### Alignments

CLC Main Workbench Version 22.0.2 (Qiagen) was used to run CLUSTAL Omega using the Additional Alignments Plugin (version 22.0 build 211216-0232-248437) to align amino acid sequences. FASTA alignment files were exported and used to run a ESPRIPT secondary structure alignment with the various PDB files for ATAD2 bromodomain (PDBID: 7M98) and ATPase domains (PDBID: 8H3H)^70,71^.

### Molecular Biology

Computational sequence analysis, design of vectors, primers and PCR fragments was performed using CLC Main Workbench (Qiagen) Version 22.0.2. Oligonucleotides were synthesised by Merck. High-fidelity PCRs were conducted using Q5 DNA polymerase (NEB) according to the manufacturer’s instructions. Low-fidelity screening PCRs were conducted using Ultra Mix Red (PCR Biosystems) according to the manufacturer’s instructions. PCR amplicons were resolved in 1% agarose (Melford) TBE gels containing 1x SYBRsafe and visualised on a Chemidoc MP (BioRad). A full list of oligonucleotides and vectors are presented in **Supplemental Data 1**. Sanger sequencing to verify plasmids etc. was conducted by Genewiz. Protein samples of cells were generated by taking 2.5 × 10^7^ log phase promastigotes, lysing in 40 μl LDS (lithium dodecyl sulfate) sample buffer supplemented to 250 mM DTT and heated to 60 °C for 10 min, after cooling, 1 μl of Basemuncher (Abcam) was added and the sample incubated at 37 °C to degrade nucleic acids. Samples were separated in 4-20% gradient TGX Stain-Free SDS-PAGE Gels (BioRad) and the total protein labelled and visualised using UV light in a BioRad ChemiDoc MP. Western blotting was performed using an iBlot II (Invitrogen) and the associated PVDF cassettes, using program P0. Membranes were blocked with 5% milk protein in 1x Tris Buffered Saline Tween-20 0.05%. Primary and secondary antibodies for western blotting are listed in **Supplemental Table 1** and were detected using appropriate fluorescent channels of chemiluminescent channels of a Chemidoc MP (BioRad), if necessary using Clarity MaxWestern ECL Substrate (BioRad).

### Parasites

Leishmania mexicana (MNYC/BZ/62/M379) derived strains were grown at 25 °C in HOMEM (Gibco) supplemented with 10% (v/v) heat-inactivated foetal bovine serum (HIFCS) (Gibco) and 1% (v/v) Penicillin/Streptomycin solution (Sigma-Aldrich). Where required parasites were grown with selective antibiotics at the following concentrations: G418 (Neomycin) at 50 μgml^−1^; Hygromycin at 50 μg ml^−1^; Blasticidin S at 10 μg ml^−1^; Puromycin at 30 μg ml^−1^ (antibiotics from InvivoGen).

### CRISPR/Cas9

BDF7 null mutants were generated in a previously published CRISPR/Cas9 screen of the *L. mexicana* bromodomain proteins using a modification of an established method^29,35^. Per gene a single sgRNA was designed with EuPaGDT to target the coding DNA sequence^72^.

Thirty residue homology flanks were identified adjacent to the CDS and appended to oligonucleotides designed to amplify drug resistance markers from blasticidin drug resistance plasmids pGL2208. After amplification of the sgRNA and resistance marker the PCR mixes were pooled and precipitated using ethanol precipitation, resuspended in sterile water and added to a transfection mix with 1 × 10^7^ mid-log promastigotes. The cell line used was *L. mexicana T7/Cas9::HYG::SAT*. Transfection was performed with an Amaxa Nucleofector 4D using program FI-115 and the Unstimulated Human T-Cell Kit. The mix was resuspended in 10 ml HOMEM 20% FCS and immediately split into two 5 ml aliquots.

Following 6–18 h of recovery time the parasites were plated at 1:5, 1:50 and 1:500 dilutions in media containing the selective drug blasticidin. Endogenous tagging was performed using the pPLOT3xMyc::mNG BSD donor vector to install N-terminal tags to BDF7, preserving the3ʹ UTR for native mRNA regulation.

### Addback strains

BDF7 CDS was amplified from *L. mexicana T7/Cas9* gDNA and cloned into pRIB Phleo using HiFi Assembly (NEB). *L. mexicana T7/Cas9 Δbdf7* promastigote forms were transfected with 10-30 µg of pRIB BDF7 that had been linearised with PacI and PmeI (NEB), gel purified then ethanol precipitated^73^. Selection was performed with 10 µg ml^-1^ phleomycin. Correct integration of the vector was assessed by PCR (**Supplemental Fig. 5**) using the oligonucleotides in **Supplemental Table 1**.

### Live-cell Microscopy

To image mNeonGreen::BDF7 10^6^ mid-log promastigotes were incubated with 1 μg ml^−1^ Hoechst 3342 for 20 min at 25 °C to stain DNA, harvested by centrifugation at 1200 × g for 10 min and washed twice with PBS. Cell pellets were resuspended in 40 μl CyGel (BioStatus) then 10 μl settled onto SuperFrost+ Slides (Thermo) and a cover slip applied. Cells were imaged using a Zeiss AxioObserver Inverted Microscope equipped with Colibri 7 narrow-band LED system and white LED for fluorescent and white light imaging. Cells were imaged using the x63 or x100 oil immersion DIC II Plan Apochromat objectives. Hoechst signal was imaged using the 385 nm LED and filter set 49, mNeonGreen with the 469 nm LED and filter set 38. Z-stacks were obtained using the Zen Blue software to control the system and exported as .CZI files to be processed in ImageJ(FIJI) using the Microvolution blind deconvolution module. Wavelength parameters were set for Hoechst (497 nm) and mNeonGreen (517 nm) emission, and refractive index parameters were defined for Cygel (1.37). Blind deconvolution was iterated 100 times using the scalar setting. Individual Z-planes for subpanels were exported in TIFF format.

### Immunofluorescence Microscopy

Promastigote cells were washed twice with PBS (1,400 g for 10 minutes at room temperature). About 106 cells in PBS were left to adhere on a poly-L-lysine-treated high precision coverslip (thickness No. 1.5H [0.170 mm ± 0.005 mm], MARIENFELD: cat. 0107222) for 15 minutes at 37°C. In vivo cross-link was then performed incubating the adhered cell with 1 mM disuccinimidyl suberate (DSS) in PBS for 10 minutes at 37°C. Attached parasites were fixed at room temperature with 4% paraformaldehyde in PBS for 15 minutes, followed by quenching with 0.1 M glycine in PBS (pH 7.6) for 5 minutes. After two washes with PBS, cells were permeabilized with 0.5% Triton X-100 in PBS for 15 minutes. Blocking was performed by incubating cells in blocking buffer (5% BSA, 0.01% saponin in PBS) for 1 hour at room temperature. Primary immunostaining was carried out for 1 hour at room temperature using mouse anti-1-tubulin KMX-1 antibody (Sigma-Aldrich, MAB3408; diluted 1:800 in blocking buffer), and rat anti-HA-Tag (clone 7C9, chromotek 7c9-100; diluted 1:200 in blocking buffer). After three washes with 0.1% Triton X-100 in PBS, cells were incubated for 1 hour with Alexa Fluor™ 568-conjugated goat anti-mouse IgG (Invitrogen A-11031 diluted 1:800) and Alexa FluorTM 647-conjugated donkey anti-rat IgG (Invitrogen A-78947; diluted 1:200) secondary antibodies prepared in blocking buffer. Following three washes with 0.1% Triton X-100/PBS, cells were counterstained with 20 µg mL-1 DAPI in PBS for 30 minutes, followed by a final PBS wash. Coverslips were mounted on glass slides using ProLong diamond antifade mountant (Invitrogen™), according to the manufacturer’s instructions.

### Microscopy and Image Analysis

Cells were examined by super-resolution structured illumination microscopy (SR-SIM), performed on a Zeiss Elyra 7 system operating in Lattice SIM2 mode, achieving an XY resolution of approximately 60 nm. Image acquisition was performed in z-stack mode, with 50 slices captured at 0.091 µm intervals. SIM reconstruction was conducted following chromatic aberration correction, via the channel alignment function in ZEN software (Zeiss), and performing deconvolution using default parameters appropriate for different fluorescent signal intensities: fixed standard for β-tubulin_AF568, fixed weak for BDF7_AF647; and low contrast for DAPI

### XL-BioID Sample Preparation

The cell lines *L. mexicana T7/Cas9 3xMyc::BirA*::BDF7*, *3xMyc::BirA*::CRK9* were generated using Cas9 directed endogenous tagging with pPlot BirA* Puro^35^ . Cultures in 100 ml HOMEM 10% FBS were set up in quadruplicate and grown until early log stage, approximately 3x10^6^ cells ml^-1^, at this point d-biotin was supplemented to the media at 150 µM for 18 hours at 25 °C. After biotinylation cells were harvested by centrifugation (1500 x *g* for 10 minutes) washed twice in PBS then resuspended in pre-warmed PBS at a density of 4 x 10^7^ cells ml^-1^. Cells were treated with 1 mM DSP crosslinker (Thermo) for 10 minutes at 25 °C; this crosslinking was quenched with Tris-HCl pH7.5 to a concentration of 20 mM. The cells were then collected by centrifugation and stored at -80 °C until lysis. The frozen cell pellets were processed as previously with the following modifications. Samples were lysed with 500 µl ice cold RIPA buffer (25 mM Tris-HCl pH 7.6, 150 mM NaCl, 1% NP-40, 1% sodium deoxycholate, 0.1% SDS) containing 2x HALT protease inhibitor cocktail (Thermo) and 1x PhosSTOP (Roche). To each tube of lysate, 1µl of BaseMuncher Endonuclease (250 units, Abcam) was added and nucleic acids digested at room temperature for 10 minutes. The samples were then sonicated using a BioRuptor Pico (3x cycles, 30 seconds on, 30 seconds off, 4 °C) and clarified by centrifugation in Protein LoBind tubes (Eppendorf) at 10, 000 x g for 10 minutes at 4 °C. Biotinylated proteins were then enriched using 100 μl of magnetic streptavidin bead suspension (1 mg of beads, ResynBioscience) for each affinity purification from 4 × 10^8^ parasites. Binding was performed overnight at 4 °C with end-over-end rotation. The next morning, beads were washed in 500 μl of the following buffers for 5 min each: RIPA for 4 x washes, 4 M urea in 50 mM triethyl ammonium bicarbonate (TEAB) pH8.5, 6 M urea in 50 mM TEAB pH8.5, 1 M KCl, 50 mM TEAB pH8.5. Beads from each affinity purification were then resuspended in 200 μl 50 mM TEAB pH8.5 containing 0.01% ProteaseMAX (Promega), 10 mM TCEP, 10 mM Iodoacetamide, 1 mM CaCl_2_ and 500 ng Trypsin Lys-C (Promega). Digest was conducted overnight in a 37 °C shaking heat block, at 900 rpm. The treatment with reducing agent in this step also cleaves the DSP crosslinker. The supernatant was recovered from the beads which were then washed with 50 µl water for 5 mins to maximise recovery of peptides.

Digests were acidified with trifluoroacetic acid (TFA) to a final concentration of 0.5% and centrifuged for 10 min at 17,000 g to remove insoluble material. The digested peptides were desalted using Strata C18-E columns (55 µm, 70 Å, 50 mg, 1ml tubes – Phenomenex), elution volume was 3 x 90 µl acetonitrile, peptides were dried down using a miVac Centrifugal Concentrator (Barnstead).

### Mass-spectrometry Acquisition

Samples were loaded onto a nanoAcquity UPLC system (Waters) equipped with a PharmaFluidics μPAC C_18_, Trapping column and a PharmaFluidics 50 cm μPAC C_18_ nano-LC column (5 μm pillar diameter, 2.5 μM inter-pillar distance). The trap wash solvent was 0.1% (v/v) aqueous formic acid and the trapping flow rate was 10 µL/min. The trap was washed for 5 min before switching flow to the capillary column. Separation used a gradient elution of two solvents (solvent A: aqueous 0.1% (v/v) formic acid; solvent B: acetonitrile containing 0.1% (v/v) formic acid). The analytical flow rate was 1 μL/min and the column temperature was 50°C. The gradient profile was linear 2.5-30% B over 30 mins then linear 30-90% B over 5 mins. All runs then proceeded to wash with 90% solvent B for 5 min. The column was returned to initial conditions and re-equilibrated for 5 min before subsequent injections.

The nanoLC system was interfaced with a maXis HD LC-MS/MS system (Bruker Daltonics) with CaptiveSpray ionisation source (Bruker Daltonics). Positive ESI-MS and MS/MS spectra were acquired using MRM mode to define data independent acquisition (DIA) windows with a width of 5 Th between *m/z* 450-650. Instrument control, data acquisition and processing were performed using Compass 1.7 software (microTOF control, Hystar and DataAnalysis, Bruker Daltonics). Instrument settings were: ion spray voltage: 1,450 V, dry gas: 3 L/min, dry gas temperature 150°C, ion acquisition range: *m/z* 280-1,600, spectra rate: 15 Hz, quadrupole low mass: 322 *m/z*, collision RF: 1,400 Vpp, transfer time 120 ms. The collision energy was set to 22 for DIA windows below *m/z* 600 and to 24 for higher *m/z* windows.

LC-MS data, in Bruker .d format, were converted to .mzML format using MSConvert (ProteoWizard) before analysing using DIA-NN (1.8.1) with searching against and in-silico predicted spectral library, derived from the LmexCas9T7-prot database appended with common proteomic contaminants. Search criteria were set to maintain a false discovery rate (FDR) of 1%. Peptide-centric output in .tsv format, was pivoted to protein-centric summaries using KNIME and data filtered to require protein q-values < 0.01 and a minimum of two peptides per accepted protein.

### Mass-spectrometry Proximal Protein Analysis

The quantification values from DIA-NN were analysed using limma in R as previously72 . Missing values were imputed by values drawn from a left-shifted normal log2 intensity distribution to model low abundance proteins (mean = 14, sd = 1.2). Proximal proteins were determined with the limma package using options trend = TRUE and robust = TRUE for the eBayes function. Multiple testing correction was carried out according to Benjamini & Hochberg, the false discovery rate for identified proximals was 1%. Radial plots were generate in R-Studio using ggPlot ^74^.

### RNA-seq

Cultures of *L. mexicana T7/Cas9*, *Δbdf7* and *Δbdf7::RIB::BDF7* were set up in Grace’s medium at 2x10^5^ cells ml^-1^ and grown at 25 °C so as to be able to recover 2x10^7^ cells at days 3, 7 and 8. After harvest on day 7 cultures were transferred to a 35 °C incubator for the final 24 hours to stimulate amastigote differentiation. RNA was extracted with the Monarch RNA miniprep kit (NEB) using the gDNA removal column, the included on-column DNAse treatment and then, subsequent to elution, the RNA was treated with the TurboDNA Free kit (Thermo) to remove all gDNA contamination. Samples were then frozen at -80 °C and transferred to Novogene on dry ice. The following methods description is provided from Novogene: A total amount of 1 µg RNA per sample was used as input material for the RNA sample preparations. Sequencing libraries were generated using NEBNext® Ultra TM RNA Library Prep Kit for Illumina® (NEB, USA) following manufacturer’s recommendations and index codes were added to attribute sequences to each sample. Briefly, mRNA was purified from total RNA using poly-T oligo-attached magnetic beads. Fragmentation was carried out using divalent cations under elevated temperature in NEBNext First Strand Synthesis Reaction Buffer (5X). First strand cDNA was synthesized using random hexamer primer and M-MuLV Reverse Transcriptase (RNase H-). Second strand cDNA synthesis was subsequently performed using DNA Polymerase I and RNase H. Remaining overhangs were converted into blunt ends via exonuclease/polymerase activities. After adenylation of 3’ ends of DNA fragments, NEBNext Adaptor with hairpin loop structure were ligated to prepare for hybridization. In order to ,select cDNA fragments of preferentially 150∼200 bp in length, the library fragments were purified with AMPure XP system (Beckman Coulter, Beverly, USA). Then 3 µl USER Enzyme (NEB, USA) was used with size-selected, adaptor ligated cDNA at 37 °C for 15 min followed by 5 min at 95 °C before PCR. Then PCR was performed with Phusion High-Fidelity DNA polymerase, Universal PCR primers and Index (X) Primer. At last, PCR products were purified (AMPure XP system) and library quality was assessed on the Agilent Bioanalyzer 2100 system. The clustering of the index-coded samples was performed on a cBot Cluster Generation System using PE Cluster Kit cBot-HS (Illumina) according to the manufacturer’s instructions. After cluster generation, the library preparations were sequenced on an Illumina platform and paired-end reads were generated. Raw reads in FASTQ format were processed using a Novogene pipeline, reads under went QC to filter low-quality reads and adapter contamination. HISAT2 was then used to map to the *Leishmania mexicana* U1103 genome retrieved from TriTrypDB^39,75^. HTseq was used to quantify gene expression using union mode, using FPKM normalisation. Differential gene expression was conducted using DEseq2^63^ .

### ChIP-seq

ChIP-seq of BDF7 was conducted using the ChIP-it Express Enzymatic Kit (Active Motif) and the procedure from^29^ , with several modifications. Firstly, a strain of *L. mexicana T7/Cas9 10xTY::BDF7* was generated using Cas9-directed endogenous tagging. Mid-log promastigotes cultures of this strain and the parental *L. mx T7/Cas9* strain were harvested to recover 3 x 10^8^ cells per ChIP replicate. Cells were fixed with 1% formaldehyde for 5 min and quenched with 1x of the included glycine solution. Nuclei were extracted from the cells using a Dounce Homogeniser and subjected to enzymatic digest and 3 x 10 second sonication bursts using Sonics Vibra-Cell probe sonicator (set to 40% amplitude). Chromatin fragmentation was checked by agarose gel electrophoresis and quantified prior to setting up the immunoprecipitations. Five micrograms of chromatin (amount of DNA) was pre-cleared with Protein G magnetic resin, and then set up with 30 µl fresh Protein G magnetic beads as per the ChIP-it Express manual. The immunoprecipitation was performed with the addition of 2 µg anti-TY antibody (Active Motif, ChIP-grade, Mouse MAb-054-050, IgG_1_) per pulldown, and incubated at 4 °C for 2 hours. Beads were washed and the crosslinking was reversed following the manufacturer’s instructions. DNA liberated from this step was purified using ChIP-cleanup mini-columns (Zymogen), quantified using the Qubit High Sensitivity DNA assay (Qiagen) and sent to Novogene for library preparation and Illumina Sequencing. Reads were quality checked with FastQC version 11.0.5 and trimmed with cutadapt/4.2-GCCcore-11.3.0. They were then aligned to the *L. mexicana* U1103 genome ^39^ and *L. mexicana T7/Cas9* assembly^43^ using bwa-mem (BWA/0.7.17 and SAMtools/1.17 ). This was conducted using the University of York Viking cluster. MACS2^40^ was accessed via Galaxy^41^ to analyse the triplicate 10xTY::BDF7 ChIP elution samples against the LmxT7/Cas9 elution (MACS2 callpeak Galaxy Version 2.2.9.1+galaxy0). Pooled treatment and control file options were used, and the BAM files were provided, with the tool run in paired-end mode. Narrow band and broad band peak options were run using the default minimum cut-off of 5-fold enrichment to call confident peaks. Co-ordinates and corresponding read depths were extracted from the BED files and used to plot the graphs in Prism Graphpad. Data was plotted as a Circos Plot using Circa software (OMGenomics).

### Artificial Intelligence Tools and Technologies

Protein structural models were accessed through the AlphaFoldDB website (https://alphafold.ebi.ac.uk). The multimeric modelling of LmxBDF7 was conducted using AlphaFold3 accessed by the AlphaFold Server (https://alphafoldserver.com)^76^ . The validity of the models was assessed using the pLDDT for local confidence of the protomer, and the pTM and iPTM scores for the confidence of the multimers. These were visualised using UCSF ChimeraX ^77^ . This data is presented in Supplemental Figure 3. No other AI tools, for example LLMs, were used in the genesis of any other part of the manuscript.

## Acknowledgements

The York Centre of Excellence in Mass Spectrometry was created thanks to a major capital investment through Science City York, supported by Yorkshire Forward with funds from the Northern Way Initiative, and subsequent support from EPSRC (EP/K039660/1; EP/M028127/1). The Viking cluster was used during this project, which is a high-performance compute facility provided by the University of York. We are grateful for computational support from the University of York, IT Services and the Research IT team.

This work was supported by funding from GSK through the Pipeline Futures Group, and a Fellowship from a Research Council United Kingdom Global Challenges Research Fund under grant agreement A Global Network for Neglected Tropical Diseases grant number MR/P027989/1. to Nathaniel Jones. This work was part-funded by the Wellcome Trust [ref: 204829] through the Centre for Future Health (CFH) at the University of York.

## Supplemental Files

Supplemental Table 1 – Oligos, Plasmids, and Antibodies

Supplemental Table 2 – 10xTY::BDF7 MACS PEAKS over 2.5 fold enriched.

Supplemental Table 3 – XL-BioID BirA*::BDF7 Proximal Proteins – output from limma using BirA*::CRK9 as a spatial control

Supplemental Table 4 – DEseq2 output Day 3 Δbdf7 vs T7/Cas9

Supplemental Table 5 – DEseq2 output Day 3 Δbdf7::RIB:BDF7 vs T7/Cas9

Supplemental Table 6 – DEseq2 output Day 7 Δbdf7 vs T7/Cas9

Supplemental Table 7 – DEseq2 output Day 7 Δbdf7::RIB:BDF7 vs T7/Cas9

Supplemental Table 8 – DEseq2 output Day 8 Δbdf7 vs T7/Cas9

Supplemental Table 9 – DEseq2 output Day 8 Δbdf7::RIB:BDF7 vs T7/Cas9

## Supporting Figures

**Supplemental Figure 1:**
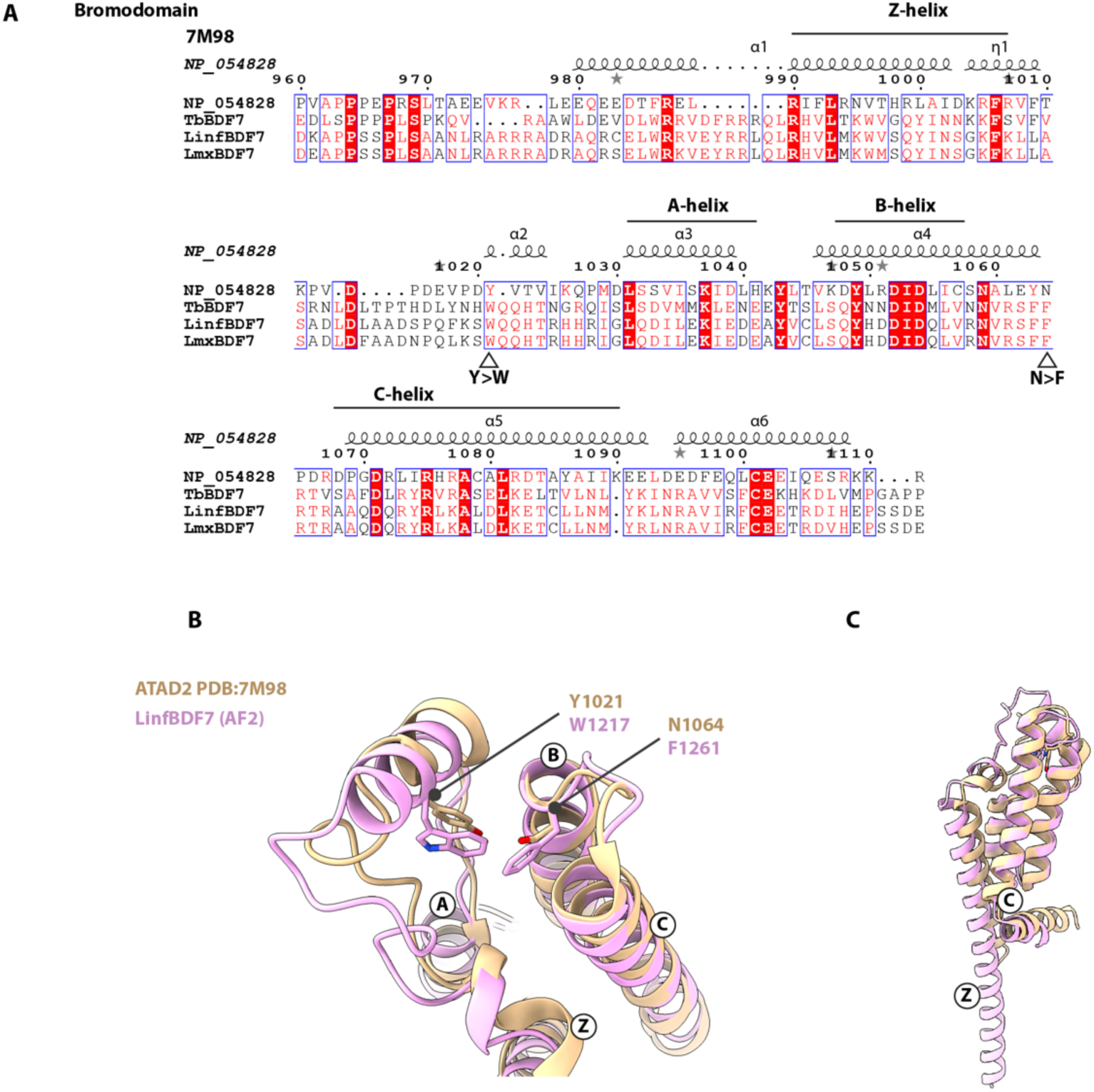
Comparison of ATAD2 bromodomain structure to BDF7 bromodomain. A. Amino acid sequence alignment of the BDF7 bromodomain with ATAD2 related bromodomains. NP_054828 is Human ATAD2, 7M98 is the PDB ID for the ATAD2 structure used to assign the secondary structures in the top line of the alignment. *T. brucei* BDF7 is included, as is *L. infantum* and *L. mexicana* BDF7. The *L. mexicana* BDF7^1143–1311^ sequence used in the alignment has 20.3% identity to the human ATAD2^960-1111^ sequence used, and 37.3% similarity. The four main helices of the bromodomain fold are annotated above the alignment, and the mutations to the conserved amino acids highlighted by open arrowheads below the sequences. B. View of the domain, emphasising the extended Z-helix and the kinked C-helix. ATAD2 possesses a bromodomain with an atypical Z-A loop structure, and contains an additional kinked, helical extension of the C-helix. These features are replicated in the AlphaFold model of *Leishmania* BDF7^6^ (AlphaFoldDB - AF-A4HV00-F1-v4) with BDF7 additionally predicted to have an N-terminal extension to the Z-helix of the bromodomain. CLUSTAL Omega^70^ alignment of the ATAD2 and Lmx BDF7 amino acid sequences, and ChimeraX Matchmaker^77^ structural alignment of an *L. infantum* BDF7 AlphaFold2 model to ATAD2 (PDBID: 7M98) were conducted to compare the two proteins. Human ATAD2 has a tyrosine at residue 1021 and an asparagine at residue 1064, consistent with the conserved residues that can co-ordinate peptide binding into the bromodomain^19^. It is evident that the Y and N residues are replaced by tryptophan 1218 and phenylalanine 1262 respectively in *Leishmania* BDF7 (Fig. 1).

**Supplemental Figure 2:**
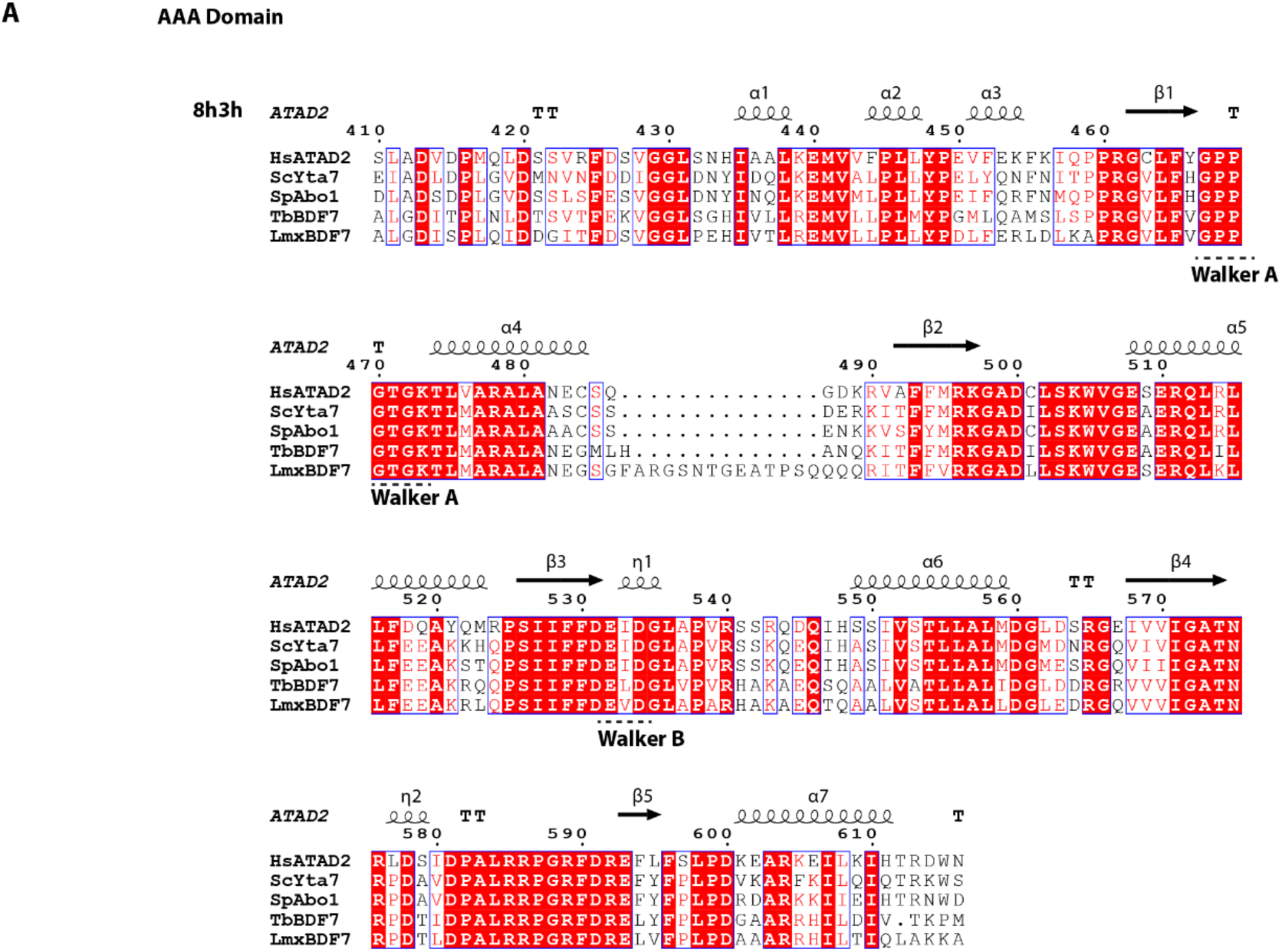
Amino acid sequence alignment of the BDF7 ATPase domain with ATAD2 related proteins. **A.** Clustal Omega alignments (as .aln files) generated in CLC Main Workbench were uploaded to ESPRIPT3 along with the PDB file for human ATAD2 bromodomain (PDB ID: 8h3h) to show the secondary structure of the aligned regions based on the 8h3h template. The Walker A and Walker B motifs are highlighted with dashed lines under the alignment.

**Supplemental Figure 3:**
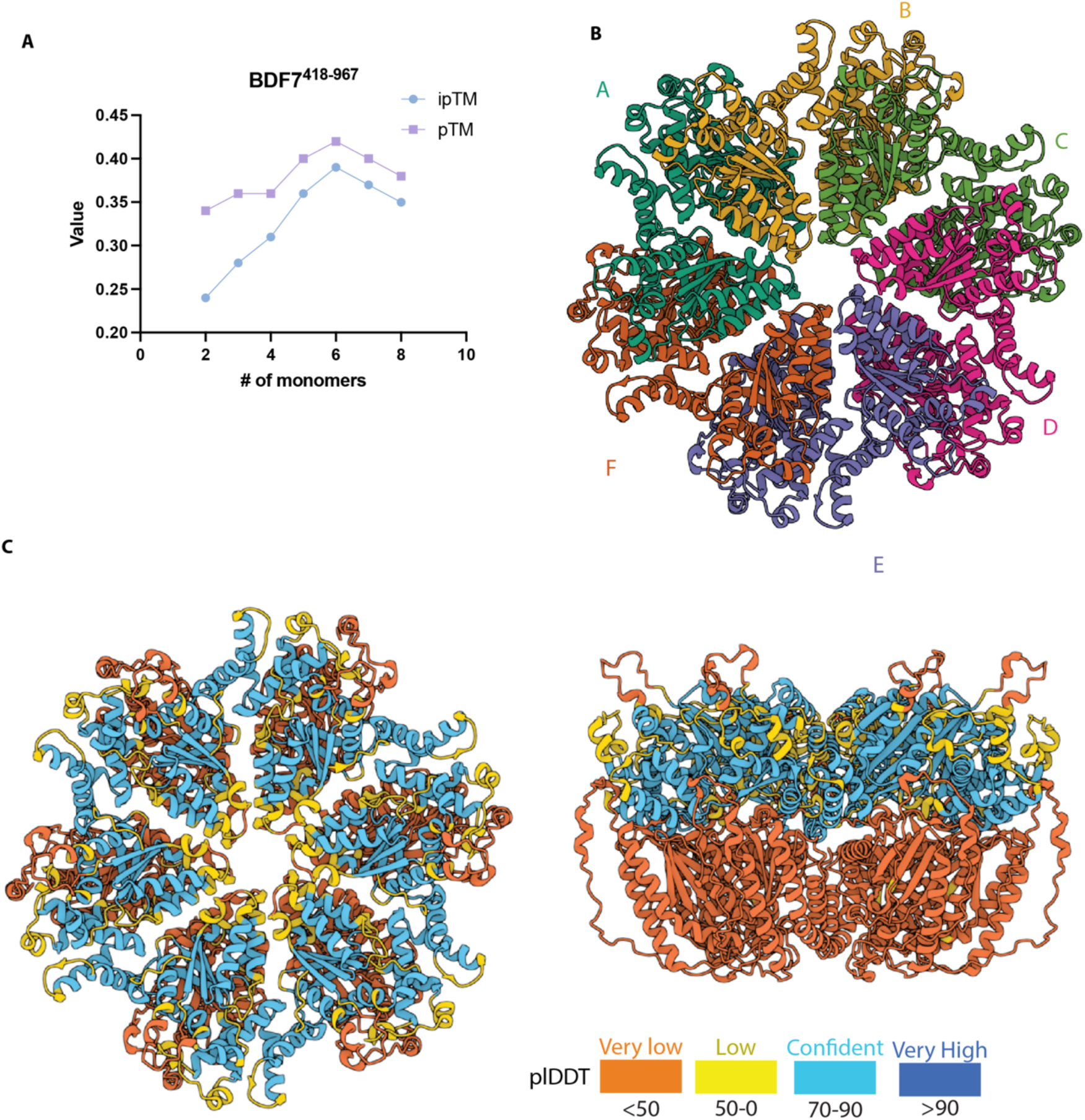
AlphaFold3 modelling of BDF7 AAA, AAA-lid and AAA-like domain. A. Line-Chart showing the changes in iPTM and pTM values as number of BDF7^418-967^ monomers are increased. B. View down onto a model of 6 units of BDF7^418-967^, with each chain coloured separately. C. The same view but with the chains coloured by plDDT scores. D. The model is rotated 90 degrees towards the viewer showing the stacking of the AAA1 domain on top of the low confidence prediction of the AAA2 domain. Due to the large size of BDF7 (1549 amino acids) and its potential to form a hexamer it was quite challenging to assess the structure with modelling tools such as AlphaFold. Using residues 418-967, which encompass the first AAA domain, the AAA lid domain and the second AAA-like domain, AlphaFold3 modelling was performed to shed light on the potential for BDF7 to exist as a multimer – as has been shown for ATAD2 and Yta7. The high plDDT scores for the subunits gives confidence (between 70-90) for the predicted structure of the AAA1 and AAA-Lid domains, in contrast to the low (<50) score for the second AAA domain – likely reflecting divergent evolution of an inactive region. Increasing the number of protomers in the model from 2-8 showed that the ipTM and pTM scores increased with each additional protomer, peaking when 6 were present, before starting to decrease. The peak values were 0.39 and 0.42 for ipTM and pTM respectively – these are both below the threshold for confident assignment of quaternary structure; however, the fact that many other AAA+ ATPases are hexameric permanently, or at some point in their reaction cycle would support the AF3 optimum. Also modelled with poor confidence was the central pore of the ATPase, potentially indicating divergence or flexibility of this region – the effects of this on substrate specificity are unknown.

**Supplemental Figure 4:**
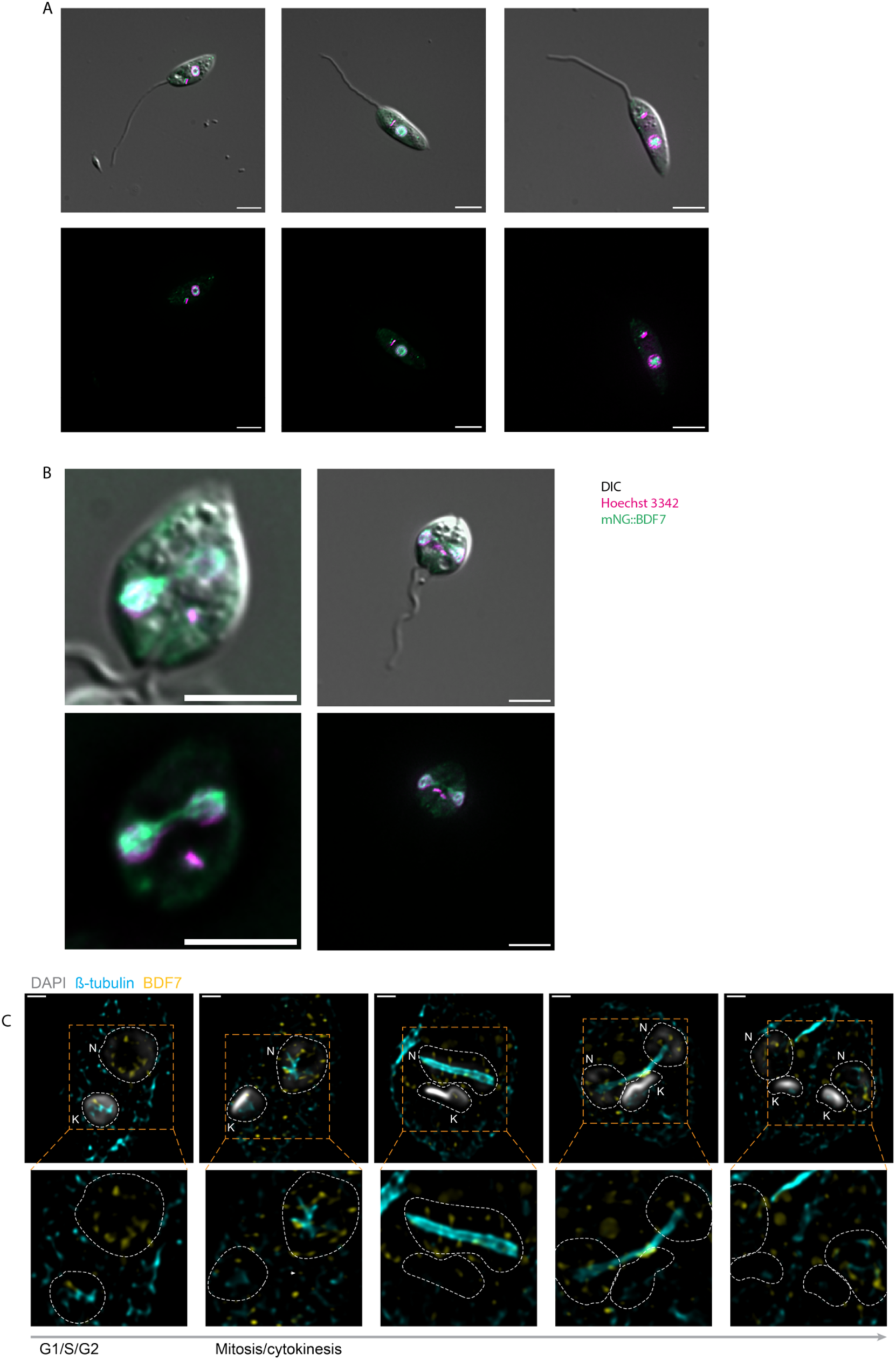
Live-cell, widefield fluorescence microscopy of Lmx T7/Cas9 mNG::BDF7. Promastigote cells were immobilised in CyGel imaged using a Zeiss AxioObserver microscope. A. Cells in G1 (1N1K) status. B. 2N2K cells exhibiting BDF7 localisation in spindle-like structures. DNA is stained with Hoechst 3342 and pseudo-coloured magenta, mNG::BDF7 is pseudo-coloured in green. DIC is Nomarsky differential interference contrast. In each image the scale bar indicates 5 µm. C. Super-resolution microscopy of immunofluorescence labelled *Lmx* 3xHA::BDF7 usingv Zeiss Structured Illumination Microscopy Elyra system. DNA is labelled with DAPI (white) and the nucleus and kinetoplasts are then outlined in dotted lines (in lower panels). β-tubulin staining (cyan) marks the mitotic spindle, BDF7 immunofluorescence signal is pseudocoloured yellow. Scale bars represent 1 µm. Series indicates a representative series of cells progressing through mitosis and cytokinesis.

**Supplemental Figure 5:**
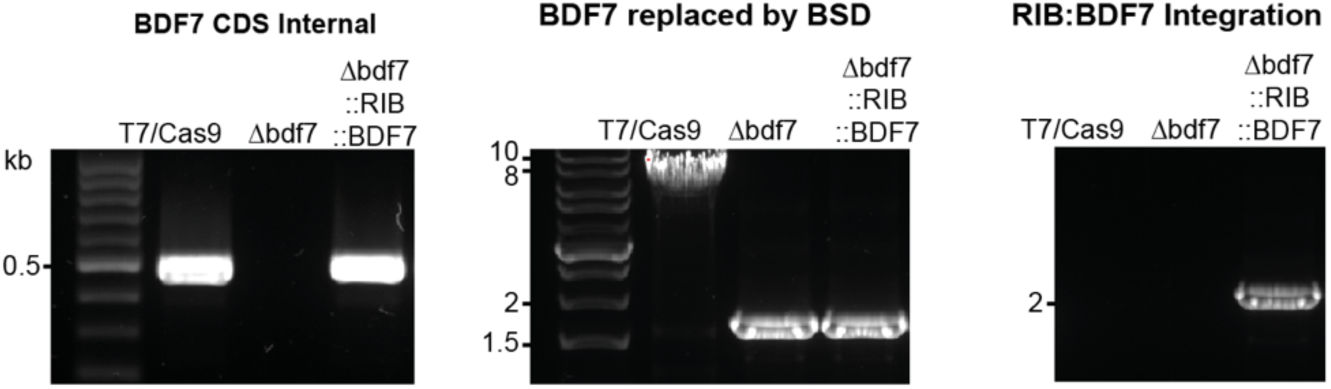
PCR validation of *BDF7* addback into Δ*bdf7* strain. Agarose gel electrophoresis of PCRs reactions to detect a *BDF7* coding sequence (left panel, 500 bp), the endogenous *BDF7* locus replaced by *BSD* (size shift ∼ 9 kb to 1.6 kb, central panel), and a successful integration of the *BDF7* expression cassette into the *RIB* locus (right panel, 2 kb).

**Supplemental Figure 6:**
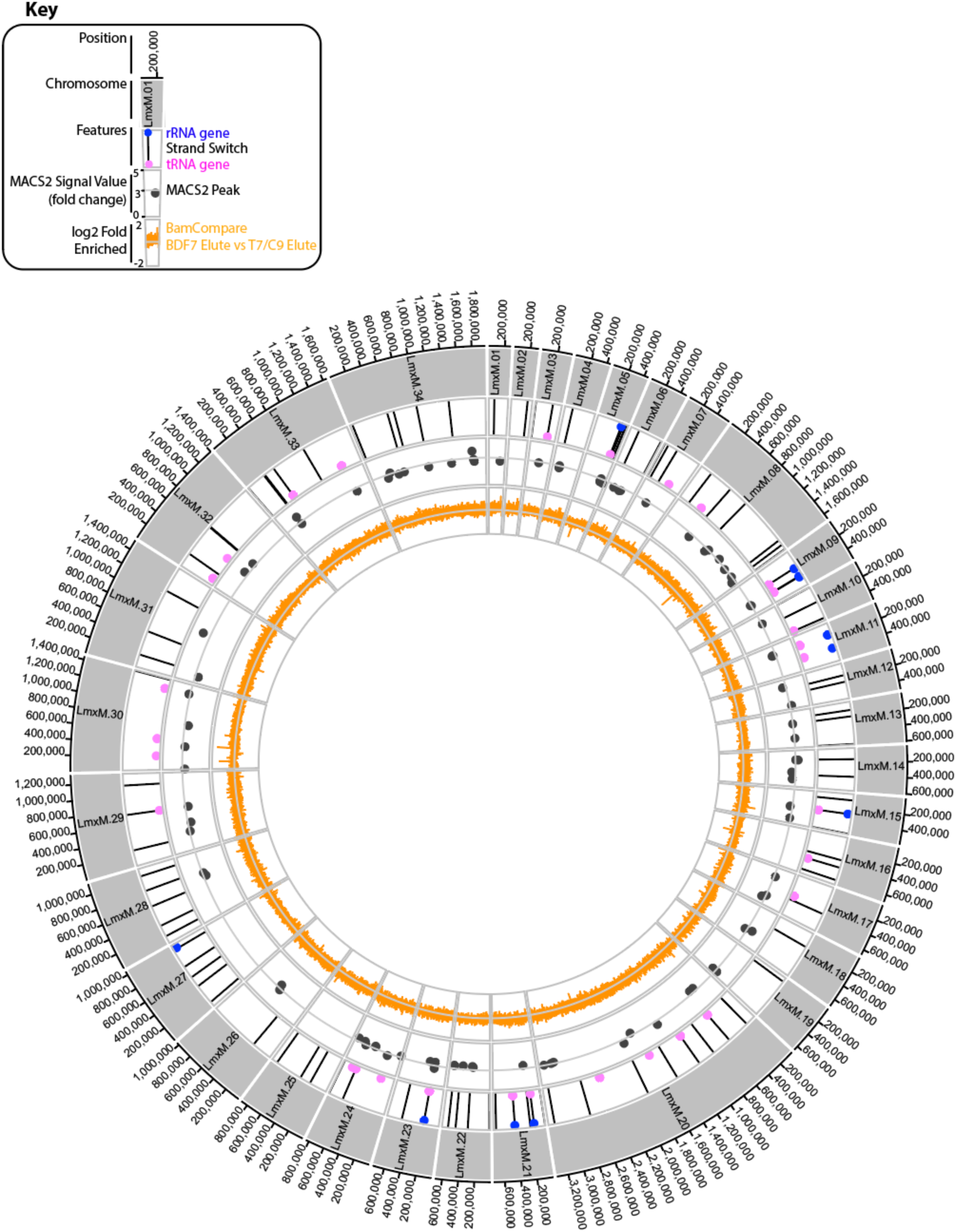
Circos Plot overview of BDF7 ChIP Data. The 34 *L. mexicana* chromosomes are depicted in the circular arrangement, with strand switch, tRNA and rRNA features in the next ring. Interior to this is the MACS2 peaks that are called, albeit under the 5-fold enrichment threshold, these were manually cut-off to show peaks giving over 2.5 fold enrichment. The grey horizontal line shows 3-fold level. The innermost ring shows the log2-fold changes when the BDF7 Elution was compared to the T7/Cas9 elution using deepTools. The grey horizontal line indicated the level of 0-fold change.

**Supplemental Figure 7:**
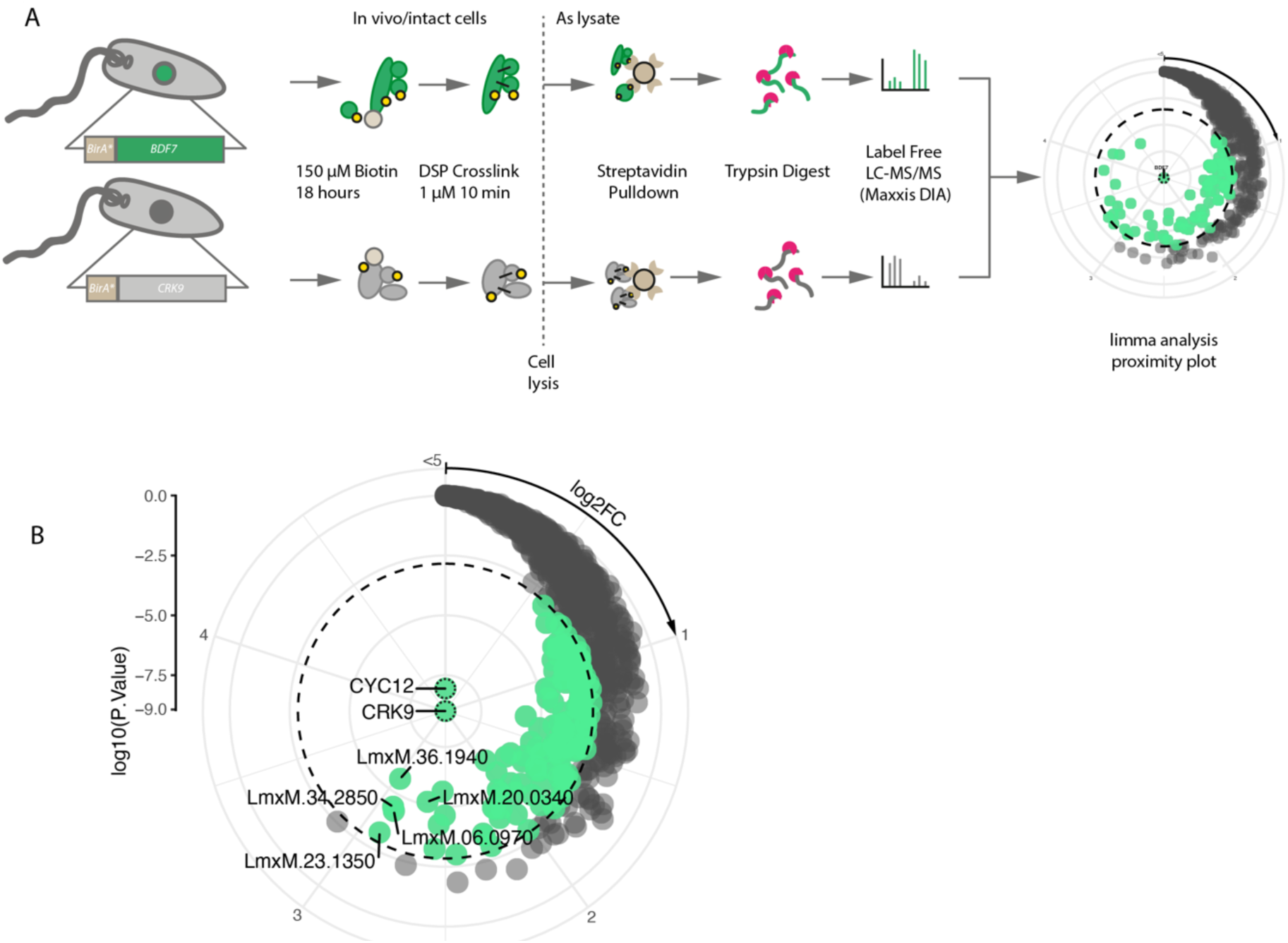
XL-BioID experimental workflow. **A**. Cartoon representing the steps taken in the XL-BioID workflow. B. Radial plot of CRK9 proximal proteins. The log2fold change is depicted on the clockwise circumferential axis, with log10(p-Value) depicted on the radial axis. Significant hits (> 2-fold , p-adj<0.01) are coloured mint green.

**Supplemental Figure 8:**
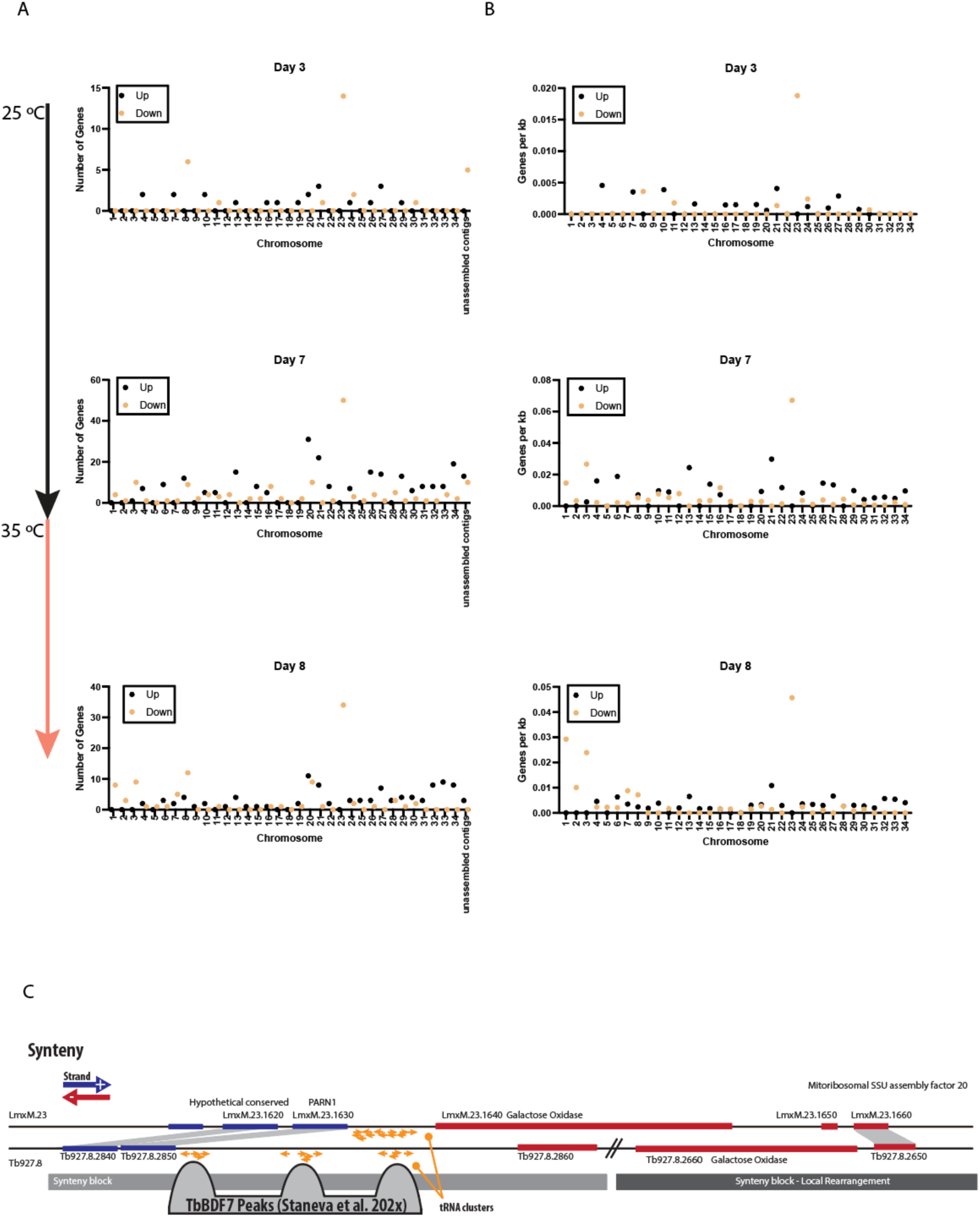
Chr 23 contains more downregulated genes than any other chromosome in the Δbdf7 strain. A. Plot showing the number of differentially expressed genes per chromosome, split by those upregulated (black points) and down regulated (tan points). B. Plot showing the number of differentially expressed genes normalised by the size of the chromosome (expressed as genes per kb) split by those upregulated (black points) and down regulated (tan points). C. Graphic representation of the conserved tRNA cluster on *Leishmania* Chr23 and T. brucei Chr8. depicting the synteny blocks and the syntenic genes in each species. There are local rearrangements in the *T. brucei* Chr8 downstream of the tRNA cluster but the synteny is maintained. Staneva et al detected mCherry::BDF7 enrichment with the tRNA clusters in T. brucei bloodstream forms. Downstream of this there are 1-2 well positioned nucleosomes in all T. brucei forms, and H2AZ. The chromatin environment at this location may represent an insulator. Coding sequences are indicated in coloured bars, blue for the + strand and red for the – strand. tRNA or ncRNA genes are indicated in orange arrows with the direction of the arrowhead indicating the strand (right for +strand, left for -strand).

**Supplemental Figure 9:**
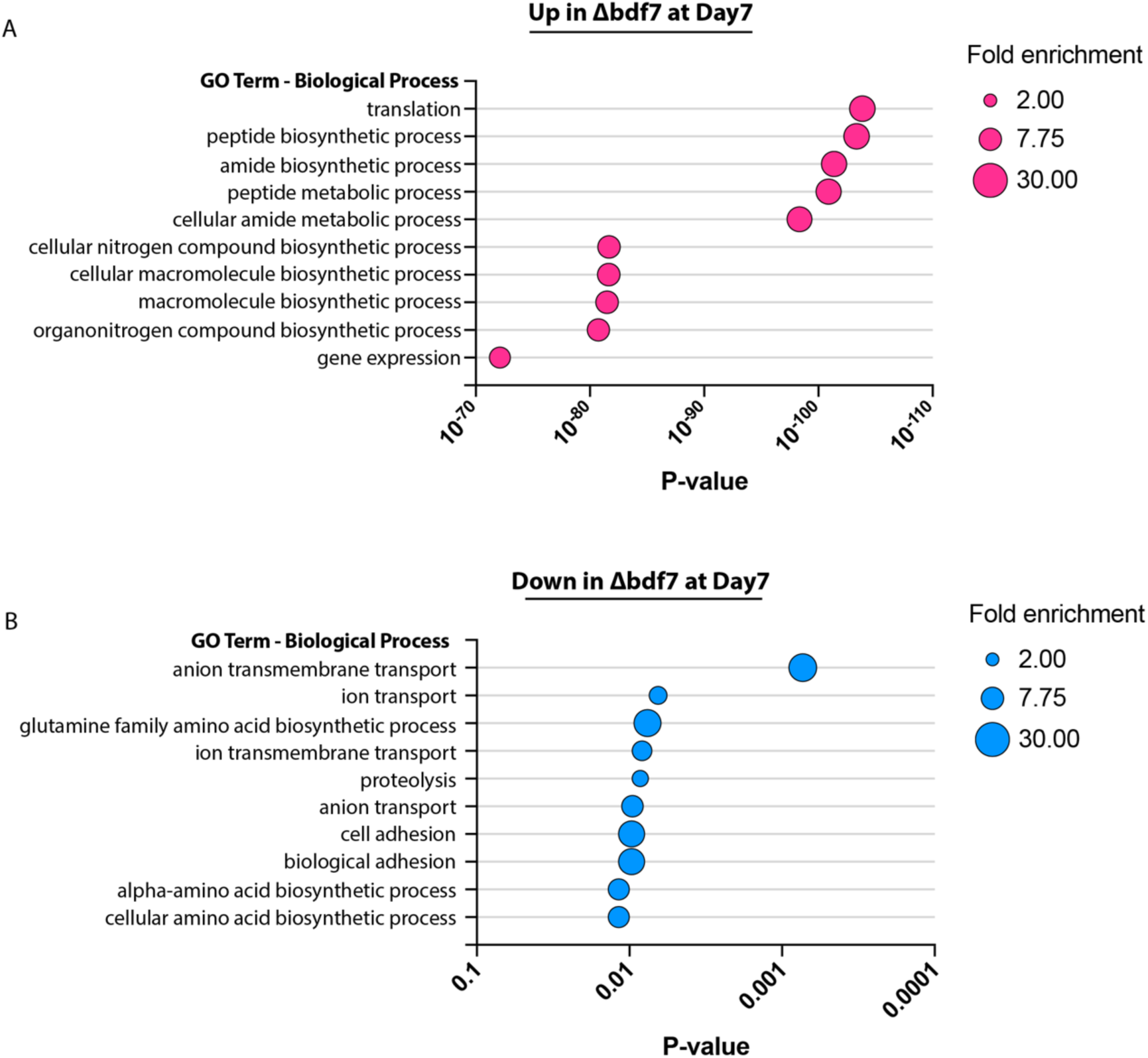
The top 10 GO terms associated with differentially expressed mRNAs in the Δbdf7 mutant (vs T7/Cas9) at day 3 in Grace’s Medium (25 °C). A. for upregulated mRNAs. B. For downregulated mRNAs. In both plots, the category of GO Term Biological Process are on the Y-axis, with the P-value on the X- axis, the size of the circle reflects the fold enrichment of the GO term in the analysis. GO term enrichment analysis was conducted with TriTrypDB Analyse Results function using the gene ID list as an input.

**Supplemental Figure 10:**
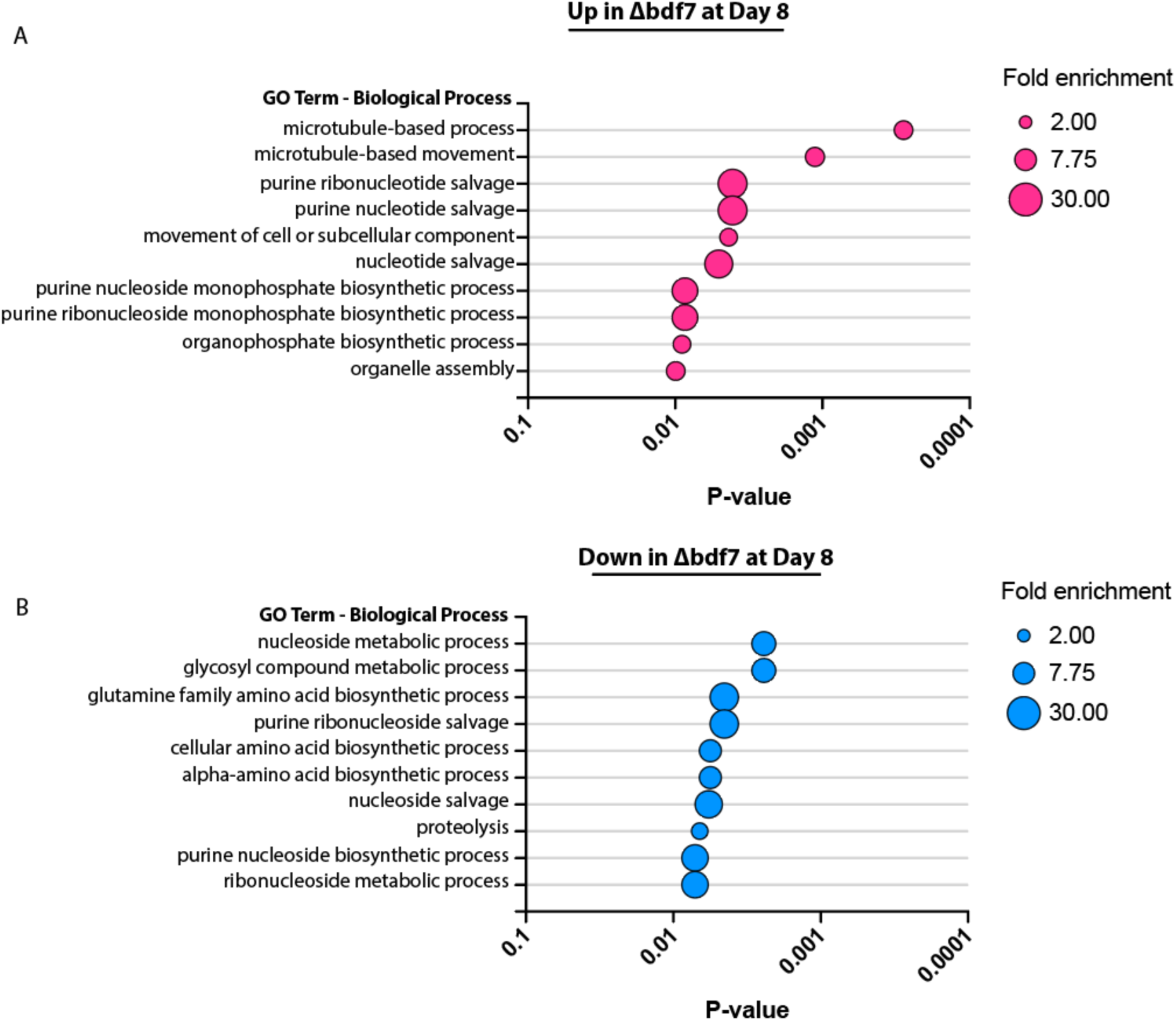
The top 10 GO terms associated with differentially expressed mRNAs in the Δbdf7 mutant (vs T7/Cas9) at day 8 in Grace’s Medium (35 °C). A. for upregulated mRNAs. B. For downregulated mRNAs. In both plots, the category of GO Term Biological Process are on the Y-axis, with the P-value on the X- axis, the size of the circle reflects the fold enrichment of the GO term in the analysis. GO term enrichment analysis was conducted with TriTrypDB Analyse Results function using the gene ID list as an input.

## Notes

### Competing Interest Statement

Felix Calderon & Raquel Gabarro are employees of GlaxoSmithKline. This work was supported by funding from GSK through the Pipeline Futures Group, and a Fellowship from a Research Council United Kingdom Global Challenges Research Fund under grant agreement A Global Network for Neglected Tropical Diseases grant number MR/P027989/1. to Nathaniel Jones. This work was part-funded by the Wellcome Trust [ref: 204829] through the Centre for Future Health (CFH) at the University of York.

## References

1. Burza, S., Croft, S. L. & Boelaert, M. Leishmaniasis. Lancet 392, 951–970 (2018).

2. Rycker, M. D., Wyllie, S., Horn, D., Read, K. D. & Gilbert, I. H. Anti-trypanosomatid drug discovery: progress and challenges. Nat Rev Microbiol 1–16 (2022) doi:10.1038/s41579-022-00777-y.

3. Catta-Preta, C. M. C., Ghosh, K., Sacks, D. L. & Ferreira, T. R. Single-cell atlas of Leishmania development in sandflies reveals the heterogeneity of transmitted parasites and their role in infection. Proc. Natl. Acad. Sci. United States Am. 121, e2406776121 (2024).

4. Williamson, K. et al. A robustly rooted tree of eukaryotes reveals their excavate ancestry. Nature 640, 974–981 (2025).

5. Jewari, C. A. & Baldauf, S. L. An excavate root for the eukaryote tree of life. Sci. Adv. 9, eade4973 (2023).

6. Filippakopoulos, P. et al. Histone Recognition and Large-Scale Structural Analysis of the Human Bromodomain Family. Cell 149, 214–231 (2012).

7. Zhou, K., Gaullier, G. & Luger, K. Nucleosome structure and dynamics are coming of age. Nat. Struct. Mol. Biol. 26, 3–13 (2019).

8. Zhou, M.-M. & Cole, P. A. Targeting lysine acetylation readers and writers. Nat. Rev. Drug Discov. 24, 112–133 (2025).

9. Caron, C. et al. Functional characterization of ATAD2 as a new cancer&sol;testis factor and a predictor of poor prognosis in breast and lung cancers. Oncogene 1–11 (2019) doi:10.1038/onc.2010.259.

10. Boussouar, F., Jamshidikia, M., Morozumi, Y., Rousseaux, S. & Khochbin, S. Malignant genome reprogramming by ATAD2. Biochim. Biophys. Acta (BBA) - Gene Regul. Mech. 1829, 1010–1014 (2013).

11. Snider, J., Thibault, G. & Houry, W. A. The AAA+ superfamily of functionally diverse proteins. Genome Biol. 9, 216 (2008).

12. Cho, C., Ganser, C., Uchihashi, T., Kato, K. & Song, J.-J. Structure of the human ATAD2 AAA+ histone chaperone reveals mechanism of regulation and inter-subunit communication. *Commun*. Biol. 6, 993 (2023).

13. Cho, C. et al. Structural basis of nucleosome assembly by the Abo1 AAA+ ATPase histone chaperone. Nat Commun 10, 5764 (2019).

14. Wang, F., Feng, X., He, Q., Li, H. & Li, H. The Saccharomyces cerevisiae Yta7 ATPase hexamer contains a unique bromodomain tier that functions in nucleosome disassembly. J Biol Chem 299, 102852 (2023).

15. Morozumi, Y. et al. Atad2 is a generalist facilitator of chromatin dynamics in embryonic stem cells. Journal of molecular cell biology 8, 349–362 (2016).

16. Wang, T. et al. ATAD2 controls chromatin-bound HIRA turnover. Life Sci. Alliance 4, e202101151 (2021).

17. Liakopoulou, A. et al. ATAD2 mediates chromatin-bound histone chaperone turnover. bioRxiv 2024.10.04.616609 (2025) doi:10.1101/2024.10.04.616609.

18. Cattaneo, M. et al. Lessons from Yeast on Emerging Roles of the ATAD2 Protein Family in Gene Regulation and Genome Organization. Mol Cells 37, 851–856 (2014).

19. Evans, C. M. et al. Coordination of Di-Acetylated Histone Ligands by the ATAD2 Bromodomain. Int J Mol Sci 22, 9128 (2021).

20. Bansal, P. & Kurat, C. F. Yta7, a chromatin segregase regulated by the cell cycle machinery. Mol. Cell. Oncol. 9, 2039577 (2022).

21. Gradolatto, A. et al. Saccharomyces cerevisiae Yta7 Regulates Histone Gene Expression. Genetics 179, 291–304 (2008).

22. Gradolatto, A. et al. A Noncanonical Bromodomain in the AAA ATPase Protein Yta7 Directs Chromosomal Positioning and Barrier Chromatin Activity. Mol Cell Biol 29, 4604–4611 (2009).

23. Kurat, C. F. et al. Restriction of histone gene transcription to S phase by phosphorylation of a chromatin boundary protein. Genes & Development 25, 2489–2501 (2011).

24. Lombardi, L. M., Davis, M. D. & Rine, J. Maintenance of nucleosomal balance in cis by conserved AAA-ATPase Yta7. Genetics 199, 105–116 (2015).

25. Lombardi, L. M., Ellahi, A. & Rine, J. Direct regulation of nucleosome density by the conserved AAA-ATPase Yta7. Proceedings of the National Academy of Sciences of the United States of America 108, E1302–11 (2011).

26. Chacin, E. et al. A CDK-regulated chromatin segregase promoting chromosome replication. Nat. Commun. 12, 5224 (2021).

27. Shahnejat-Bushehri, S. & Ehrenhofer-Murray, A. E. The ATAD2/ANCCA homolog Yta7 cooperates with Scm3HJURP to deposit Cse4CENP-A at the centromere in yeast. Proc. Natl. Acad. Sci. 117, 5386–5393 (2020).

28. Clayton, C. Regulation of gene expression in trypanosomatids: living with polycistronic transcription. Open biology 9, 190072–24 (2019).

29. Jones, N. G. et al. Bromodomain factor 5 is an essential regulator of transcription in Leishmania. Nat Commun 13, 4071 (2022).

30. Russell, C. N. et al. Bromodomain Factor 5 as a Target for Antileishmanial Drug Discovery. ACS Infect. Dis. 9, 2340–2357 (2023).

31. Tallant, C. et al. Expanding Bromodomain Targeting into Neglected Parasitic Diseases. Acs Infect Dis 7, 2953–2958 (2021).

32. Chen, F., Mackey, A. J., Stoeckert, C. J. & Roos, D. S. OrthoMCL-DB: querying a comprehensive multi-species collection of ortholog groups. Nucleic Acids Res. 34, D363–D368 (2006).

33. Staneva, D. P. et al. A systematic analysis of Trypanosoma brucei chromatin factors identifies novel protein interaction networks associated with sites of transcription initiation and termination. Genome Res 31, gr.275368.121 (2021).

34. Goos, C., Dejung, M., Janzen, C. J., Butter, F. & Kramer, S. The nuclear proteome of Trypanosoma brucei. PLoS ONE 12, e0181884 (2017).

35. Beneke, T. et al. A CRISPR Cas9 high-throughput genome editing toolkit for kinetoplastids. Royal Society Open Science **May**, (2017).

36. Halliday, C. et al. Cellular landmarks of Trypanosoma brucei and Leishmania mexicana. Mol. Biochem. Parasitol. 230, 24–36 (2019).

37. Carnielli, J. B. T. et al. The mitotic spindle kinase MSK co-ordinates segregation of the nucleus and kinetoplast in Leishmania mexicana. bioRxiv 2025.09.19.677349 (2025) doi:10.1101/2025.09.19.677349.

38. Brinkman, A. B. & Stunnenberg, H. G. Epigenomics. 3–18 (2009) doi:10.1007/978-1-4020-9187-2_1.

39. Rogers, M. B. et al. Chromosome and gene copy number variation allow major structural change between species and strains of Leishmania. Genome research 21, 2129–2142 (2011).

40. Zhang, Y. et al. Model-based Analysis of ChIP-Seq (MACS). Genome Biol 9, R137 (2008).

41. Community, T. G. et al. The Galaxy platform for accessible, reproducible, and collaborative data analyses: 2024 update. Nucleic Acids Res. 52, W83–W94 (2024).

42. Ramírez, F. et al. deepTools2: a next generation web server for deep-sequencing data analysis. Nucleic Acids Res 44, W160–W165 (2016).

43. Beneke, T. et al. Genome sequence of Leishmania mexicana MNYC/BZ/62/M379 expressing Cas9 and T7 RNA polymerase. Wellcome Open Res. 7, 294 (2022).

44. Hitchcock, R. A., Thomas, S., Campbell, D. A. & Sturm, N. R. The promoter and transcribed regions of the Leishmania tarentolae spliced leader RNA gene array are devoid of nucleosomes. BMC Microbiol. 7, 44–44 (2007).

45. McDonald, J. R. et al. Localization of Epigenetic Markers in Leishmania Chromatin. Pathogens 11, 930 (2022).

46. Garcia-Silva, M. et al. Identification of the centromeres of Leishmania major: revealing the hidden pieces. Embo Rep 18, 1968–1977 (2017).

47. Geoghegan, V., Mottram, J. C. & Jones, N. G. Tag Thy Neighbour: Nanometre-Scale Insights Into Kinetoplastid Parasites With Proximity Dependent Biotinylation. Front Cell Infect Mi 12, 894213 (2022).

48. Geoghegan, V., Jones, N. G., Dowle, A. & Mottram, J. C. Protein kinase signalling at the Leishmania kinetochore captured by XL-BioID. bioRxiv (2021) doi:10.1101/2021.07.08.451598.

49. Gosavi, U., Srivastava, A., Badjatia, N. & Günzl, A. Rapid block of pre-mRNA splicing by chemical inhibition of analog-sensitive CRK9 in Trypanosoma brucei. Molecular Microbiology 113, 1225–1239 (2020).

50. Badjatia, N., Ambrósio, D. L., Lee, J. H. & Günzl, A. Trypanosome cdc2-related kinase 9 controls spliced leader RNA cap4 methylation and phosphorylation of the RNA polymerase II subunit RPB1. Molecular and cellular biology (2013) doi:10.1128/mcb.00156-13.

51. Lou, R. & Shui, W. Acquisition and Analysis of DIA-Based Proteomic Data: A Comprehensive Survey in 2023. Mol. Cell. Proteom. 23, 100712 (2024).

52. Ritchie, M. E. et al. limma powers differential expression analyses for RNA-sequencing and microarray studies. Nucleic Acids Res 43, e47–e47 (2015).

53. Carnielli, J. B. T. et al. Chemical genetics reveals Leishmania KKT2 and CRK9 kinase activity is required for cell cycle progression. bioRxiv 2025.09.08.674917 (2025) doi:10.1101/2025.09.08.674917.

54. Zhou, Q. et al. Faithful chromosome segregation in Trypanosoma brucei requires a cohort of divergent spindle-associated proteins with distinct functions. Nucleic Acids Res. 46, 8216–8231 (2018).

55. Granneman, S. et al. The human Imp3 and Imp4 proteins form a ternary complex with hMpp10, which only interacts with the U3 snoRNA in 60–80S ribonucleoprotein complexes. Nucleic Acids Res. 31, 1877–1887 (2003).

56. Liang, X. et al. Structural snapshots of human pre-60S ribosomal particles before and after nuclear export. Nat. Commun. 11, 3542 (2020).

57. Burger, F., Daugeron, M.-C. & Linder, P. Dbp10p, a putative RNA helicase from Saccharomyces cerevisiae, is required for ribosome biogenesis. Nucleic Acids Res. 28, 2315–2323 (2000).

58. Zhang, C.-J. et al. The Arabidopsis acetylated histone-binding protein BRAT1 forms a complex with BRP1 and prevents transcriptional silencing. Nature communications 7, 11715 (2016).

59. Rajan, K. S. et al. Structural and mechanistic insights into the function of Leishmania ribosome lacking a single pseudouridine modification. Cell Rep. 43, 114203 (2024).

60. Rajan, K. S. et al. A single pseudouridine on rRNA regulates ribosome structure and function in the mammalian parasite Trypanosoma brucei. Nat. Commun. 14, 7462 (2023).

61. Bates, P. A. Complete developmental cycle of Leishmania mexicana in axenic culture. Parasitology 108, 1–9 (1994).

62. Bates, P. A. Complete developmental cycle of Leishmania mexicana in axenic culture. Parasitology 108 **( Pt** **1**), 1–9 (1994).

63. Love, M. I., Huber, W. & Anders, S. Moderated estimation of fold change and dispersion for RNA-seq data with DESeq2. Genome Biol. 15, 550 (2014).

64. Marques, C. A., Dickens, N. J., Paape, D., Campbell, S. J. & Mcculloch, R. Genome-wide mapping reveals single-origin chromosome replication in Leishmania, a eukaryotic microbe. Genome Biology 1–12 (2015) doi:10.1186/s13059-015-0788-9.

65. Díaz-Viraqué, F., Ehrlich, R. & Robello, C. Genomic Organization of Trypanosoma cruzi tRNA Genes. Genome Biol. Evol. 17, evaf108 (2025).

66. Jambunathan, N. et al. Multiple bromodomain genes are involved in restricting the spread of heterochromatic silencing at the Saccharomyces cerevisiae HMR-tRNA boundary. Genetics 171, 913–922 (2005).

67. Saunders, E. C. et al. Induction of a Stringent Metabolic Response in Intracellular Stages of Leishmania mexicana Leads to Increased Dependence on Mitochondrial Metabolism. PLoS Pathog. 10, e1003888 (2014).

68. Grünebast, J., Lorenzen, S., Zummack, J. & Clos, J. Life Cycle Stage-Specific Accessibility of Leishmania donovani Chromatin at Transcription Start Regions. Msystems 6, e00628–21 (2021).

69. Grünebast, J., Lorenzen, S. & Clos, J. Genome-wide quantification of polycistronic transcription in Leishmania major. mBio 16, e0224124 (2024).

70. Sievers, F. et al. Fast, scalable generation of high-quality protein multiple sequence alignments using Clustal Omega. Mol. Syst. Biol. 7, MSB201175 (2011).

71. Robert, X. & Gouet, P. Deciphering key features in protein structures with the new ENDscript server. Nucleic Acids Res. 42, W320–W324 (2014).

72. Peng, D. & Tarleton, R. Short Paper EuPaGDT : a web tool tailored to design CRISPR guide RNAs for eukaryotic pathogens. 1–7 (2017) doi:10.1099/mgen.0.000033.

73. Mißlitz, A., Mottram, J. C., Overath, P. & Aebischer, T. Targeted integration into a rRNA locus results in uniform and high level expression of transgenes in Leishmania amastigotes. Mol. Biochem. Parasitol. 107, 251–261 (2000).

74. Wickham, H. ggplot2, Elegant Graphics for Data Analysis. (2016) doi:10.1007/978-3-319-24277-4.

75. Shanmugasundram, A., et al. TriTrypDB: An integrated functional genomics resource for kinetoplastida. PLOS Neglected Trop. Dis. 17, e0011058 (2023).

76. Abramson, J. et al. Accurate structure prediction of biomolecular interactions with AlphaFold 3. Nature 630, 493–500 (2024).

77. Meng, E. C. et al. UCSF ChimeraX : Tools for structure building and analysis. Protein Sci. 32, e4792 (2023).

